# The RNA encoding the microtubule-associated protein tau has extensive structure that affects its biology

**DOI:** 10.1101/580407

**Authors:** Jonathan L. Chen, Walter N. Moss, Adam Spencer, Peiyuan Zhang, Jessica L. Childs-Disney, Matthew D. Disney

## Abstract

Tauopathies are neurodegenerative diseases that affect millions of people worldwide including those with Alzheimer’s disease. While many efforts have focused on understanding the role of tau protein in neurodegeneration, there has been little done to systematically analyze and study the structures within tau’s encoding RNA and their connection to disease pathology. Knowledge of RNA structure can provide insights into disease mechanisms and how to affect protein production for therapeutic benefit. Using computational methods based on thermodynamic stability and evolutionary conservation, we identified structures throughout the tau pre-mRNA, especially at exon-intron junctions and within the 5′ and 3′ untranslated regions (UTRs). In particular, structures were identified at twenty exon-intron junctions. The 5′ UTR contains one structured region, which lies within a known internal ribosome entry site. The 3′ UTR contains eight structured regions, including one that contains a polyadenylation signal. A series of functional experiments were carried out to assess the effects of mutations associated with mis-regulation of alternative splicing of exon 10 and to identify regions of the 3′ UTR that contain *cis*-regulatory elements. These studies defined novel structural regions within the mRNA that affect stability and pre-mRNA splicing and may lead to new therapeutic targets for treating tau-associated diseases.

---

RNA structures function in normal cellular processes, such as splicing, protein synthesis, and regulation of gene expression.^*1*^ At the same time, mutations that disrupt RNA structure or formation of ribonucleoproteins (RNPs) can be deleterious and cause disease.^*2*^ This has generated interest in targeting RNA with therapeutics. Evolutionarily conserved RNA structures across species may have common functions.^*3*^ However, the superficial lack of sequence conservation in noncoding RNA (ncRNA) may complicate the search for conserved structures.^*4, 5*^ Thus, specialized techniques are needed for discovery of homologous structured regions in RNA.^*4, 6*^ One method to identify stable, conserved structures combines sequence alignment with thermodynamic-based folding algorithms.^*6*^

Tauopathies are a class of neurodegenerative diseases characterized by the presence of tau inclusion bodies.^*7*^ Tauopathies such as Alzheimer’s and Parkinson’s diseases (AD and PD, respectively) are burdensome socioeconomically and affect more than 35 million and 6.3 million people, respectively, worldwide.^*8*^ Currently available treatments are largely focused on symptoms and do not target underlying disease mechanisms.^*7*^ The tau protein, which binds to microtubules and promotes microtubule assembly and stability, is encoded by the microtubule associated protein tau (*MAPT*) gene on chromosome 17.^*8-11*^ The *MAPT* gene is well conserved, with 97 to 100% homology among primates.^*12*^ This 134 kb gene is comprised of 16 exons, among which exons 2, 3, and 10 are known to be alternatively spliced, generating six isoforms ranging from 352 to 441 amino acids in length.^*8, 10-12*^ Exons 2 and 3 encode for N-terminal domains while exons 9 to 12 encode microtubule binding domains (MBD).^*9*^ With such complex processing, the *MAPT* mRNA is likely rich in conserved regulatory structures that may have important functions and may be implicated in tau-associated diseases.

Tau proteins bind to and stabilize microtubules via their MBD repeat sequences that interact with negatively charged tubulin residues via their net positive charge.^*9*^ Alterations in the protein coding content of the mRNA, including the number of MBDs, are due to alternative splicing. For example, exon 10 encodes an MBD and is alternatively spliced, resulting in protein isoforms with four (4R) or three (3R) microtubule-binding domains. In normal tissues, the 3R-to-4R (repeat) ratio is approximately 1. Deregulation of exon 10 alternative splicing due to mutation manifests itself in various diseases,^*8-11*^ including frontotemporal dementia with parkinsonism-17 (FTDP-17) and Pick’s disease, which express 4R and 3R tau at aberrantly high levels, respectively.^*8*^ In FTDP-17, various mutations at the exon 10-intron 10 junction destabilize a stem-loop structure, resulting in increased interaction of the 5′ splice site with U1 snRNA and hence increased exon 10 inclusion and 4R tau levels.^*13, 14*^ These studies suggest that structures in *MAPT* pre-mRNA intron-exon junctions affect MAPT biology and that structures elsewhere in the mRNA could as well. Indeed, previous studies have shown that regions of the 3′ untranslated region (UTR) affect translation of the tau mRNA.^*15*^

Previous studies have identified conserved sequences in *MAPT* mRNA including the exon 6-intron 7 junction,*16* an intronic sequence upstream of exon 6 (180 nucleotides),^*16*^ the intron downstream of exon 2,^*17*^ and the intron downstream of exon 10, which contains several islands of conserved sequence.^*18*^ Herein, computational studies were carried out to search for potentially conserved <u>structures</u> in *MAPT* pre-mRNA. Specifically, structured regions were identified at exon-intron junctions and within the long 3′ UTR isoform that may control tau expression. This method may be applied to diseases, such as tauopathies, in which RNA dysfunction contributes to disease development. Knowledge of RNA structure in the *MAPT* gene may lead to new therapeutics against RNA, such as small molecules for treatment of tauopathies.^*19*^

## METHODS

### Structure Analysis

RNA sequences were obtained from the National Center for Biotechnology Information (NCBI). Sequences were folded using RNAfold from the ViennaRNA 2.4.0 package.^*20*^ Sequences 500 nt in length centered around each exon-intron junction were folded in 30 nt windows every 10 nt from the 5′ end, while a 4163 nt 3′ UTR sequence of the *MAPT* gene was folded in 150 nt windows every 10 nt from the 5′ end.^*15*^ For each sequence, the free energy of each window, the average free energy of a set of 50 randomized sequences for the same window, and a z-score for the difference between the native and average random free energies were calculated. Additionally, the native sequence ensemble diversity (ED) was calculated from a partition function.^*6, 21, 22*^ The z-score is a measure of the number of standard deviations (SDs) by which the free energy of the minimum free energy (MFE) structure deviates from the average free energy of structures formed by randomized sequences.^*23, 24*^ A low z-score indicates that a structure is more stable than random and may be ordered to form structure (e.g. by evolution). The *p*-value represents the fraction of sequences in the randomized sequences whose free energy is lower than the MFE of a given sequence.^*21*^ The ED is a measure of the difference (in base pairs) among Boltzmann-weighted structures in a predicted ensemble; here, a lower than average ED for a sequence indicates less conformationally diversity (e.g. a single dominant conformation).^*24, 25*^ Folded sequences whose z-scores were more than one SD below the average z-score of all windows were compiled and refolded to generate a new set of structures. Next, 250 nt sequences centered on each exon-intron junction were folded using 70 nt windows and 1 nt base step size. These parameters were chosen to identify relatively small motifs at exon-intron junctions that may be not be predicted by larger window or base step sizes.

NCBI BLASTn was used to search for sequence fragments in non-human primates from stable regions. Resulting sequence fragments were filtered to exclude duplicate fragments and fragments with less than 80% length of the query sequence. Sequences were aligned with the Multiple Alignment using Fast Fourier Transform (MAFFT) program^*26*^ and folded with RNAalifold^*20*^ to generate a consensus structure. The number of base pairs from each aligned sequence in the consensus structure was counted. The percent of canonical base pairs for each nucleotide pair and the total average percent of canonical base pairs were calculated.

Probabilities of loops in predicted and SHAPE-constrained structures of the 5′ UTR were calculated using ProbScan.^*27*^ ProbScan estimates the probabilities of loops in secondary structure according to the frequency that a particular loop appears in an ensemble of structures, which corresponds to its approximate probability in the ensemble.^*27*^ The APSI is a measure of similarity between aligned sequences.^*6, 28*^ The effects of mutations on structures at exon-intron junctions on structure were analyzed with RNA2DMut.^*29*^ Using RNAfold, RNA2DMut calculates for a given input sequence, an MFE structure and its Gibbs folding energy (ΔG°_37_) in kcal/mol.^*29*^ Additionally, it calculates a partition function, a centroid structure, defined as the structure that is most similar to other structures in the ensemble, and the ED for the input sequence.^*29*^

### Cell-based assays

Mini-gene reporters of exon 10 splicing, whether wild type tau, tau with intron 9 G(−10)T, intron 10 G(+3)A, or intron 10 C(+14)T mutations^*30*^, or fragments of a long 3′ UTR isoform^*15*^ were used in cell-based experiments. Mini-genes expressing wild type tau, tau with intron 10 C(+14)T mutations, and a long 3′ UTR isoform (4163 nt) were provided by Prof. Michael S. Wolfe (The University of Kansas).^*30-32*^ Mini-genes expressing tau with intron 9 G(−10)T, intron 10 G(+3)A, and fragments of the long 3′ UTR isoform were generated using a QuikChange Site-Directed Mutagenesis Kit (Agilent) and the manufacturer’s recommended protocol. Mini-genes containing mutations within intron 9 or 10 express a firefly luciferase reporter gene in-frame with exon 10, which leads to a decrease in signal when exon 10 is excluded (Figure 1).

**Figure 1:**
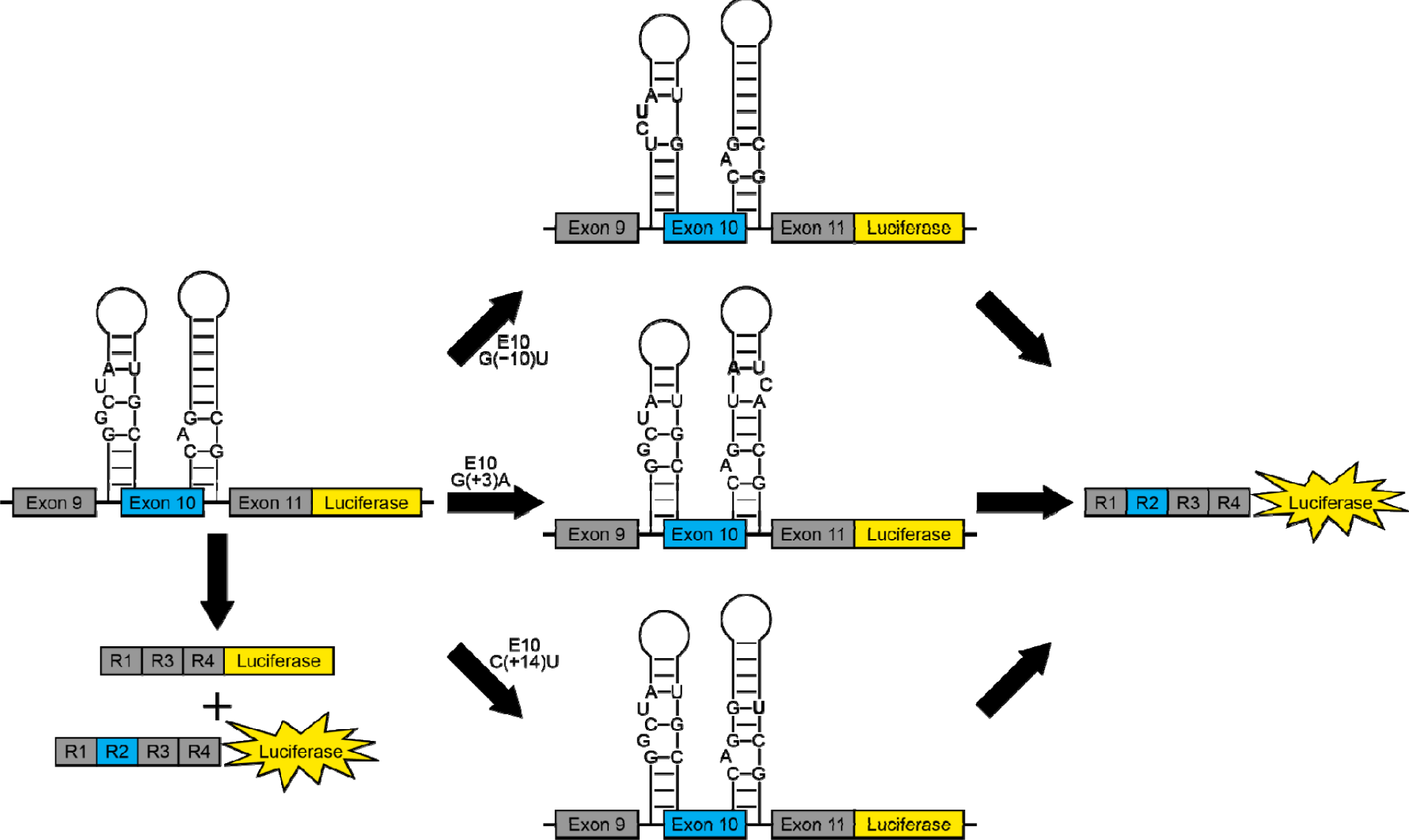
Schematic representation of cell-based assays with mini-genes expressing wild type tau or tau with mutations in intron 9 or 10.^*30, 32*^ Mini-genes contain exon 10 in frame with a firefly luciferase reporter gene. Wild type mini-genes express tau in a 4R-to-3R ratio of ∼1. Mutations in this study (indicated in bold) that affect structures at splice sites adjacent to exon 10 lead to increased exon 10 inclusion and consequently, increased luciferase signal.

HeLa cells were maintained in Dulbecco’s Modified Eagle Medium (DMEM) with 10% fetal bovine serum (FBS) and 1% GlutaMax at 37 °C. For experiments with mini-genes containing intron 9-exon 10 or exon 10-intron 10 junctions, cells were transfected at ∼90% confluency in a 100 mm dish using jetPRIME (PolyPlus-transfection SA) with 10 μg of mini-gene. A 60 mm dish was mock transfected using jetPRIME as a control. Cells were resuspended in growth medium, seeded in a 384-well plate at 10,000 cells per well, and incubated at 37 °C for 24 h. After, 5 μL of CellTiter Fluor reagent (Promega) was added to each well, the cells were incubated at 37 °C for 30 min, and fluorescence was measured (380 nm/505 nm excitation/emission wavelegnths with a 495 nm cutoff filter) using a SpectraMax M5 plate reader (Molecular Devices). Next, 25 μL of ONE-Glo reagent (Promega) was added to each well, and cells were incubated at room temperature for 3 min. Luminescence was measured on a SpectraMax M5 plate reader with a 500 ms integration time.

Mini-genes expressing fragments of a long 3′ UTR isoform were fused to firefly luciferase in the context of pmiRGLO vector, which also express Renilla luciferase used for normalization (see Supplementary Information for cloning methods). Fragments can either increase or decrease firefly luciferase signal, corresponding to increased or decreased tau expression, respectively. HeLa cells were seeded at 250,000 cells per well in 6-well plates and 2 mL of growth medium. At ∼60% confluency, cells were transfected with the mini-gene of interest, resuspended in growth medium, seeded in a 96-well plate at 50,000 cells per well density, and incubated at 37 °C. After 24 h, growth medium was removed from each well, and the cells were washed with 50 μL of 1× DPBS and lysed with 20 μL of lysis buffer. A 50 μL aliquot of firefly luciferin buffer was added to each well, and luminescence was measured as described above. Next, 50 μL of Renilla luciferase buffer was added to each well, the cells were incubated for 10 min at room temperature, and luminescence was measured as described above.

## RESULTS AND DISCUSSION

### Overview of computational analysis

The prediction of RNA secondary (2D) structure is an active area of research, and a number of algorithms attempt to find the native 2D conformation from sequence.^*33*^ Despite varying approaches, almost all folding algorithms use the thermodynamic energy of folding in prediction. The free energy change (ΔG°) going from the unfolded to the folded state of RNA 2D structure measures its energetic favorability. The ΔG° of RNA 2D structure can be calculated from the “Turner Rules,” a compilation of free energy parameters for RNA 2D structures that is based on experimental measurements of small motifs.^*34, 35*^ In the most widely applied folding algorithms (e.g. mFold^*36*^, RNAfold^*37*^ and RNAstructure^*38*^) the Turner Rules are used to find the <u>m</u>inimum free <u>e</u>nergy (MFE) 2D structure of a sequence in silico;^*34, 39*^ here the MFE (most stable) fold is assumed to best reflect the native structure.

The MFE ΔG° indicates the stability of a 2D structure, *but not its potential for function*. It was observed, however, that compared to random sequences, the MFE ΔG° of functional RNAs is lower.^*40*^ This is attributed to the *evolved order and composition* of functional RNAs, which require structure to function; disrupting this ordered sequence also disrupts stabilizing base pairs leading to less favorable ΔG° values. This propensity for folding is quantified by the ΔG° z-score 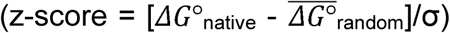. That is, ΔG° z-score measures the stability of the native sequence vs. random sequence and suggests an evolved property of the RNA.^*23, 40*^ The z-score metric is indeed at the heart of functional 2D structure prediction approaches^*41, 42*^ and was used to identify conserved functional regions in the genomes of influenza viruses^*6, 43*^ and EBV,^*21*^ as well as the mammalian Xist long <u>n</u>on<u>c</u>oding (lnc)RNA.^*44*^ These analyses led to the discovery of functional *cis*-regulatory RNA structures^*45-47*^ and novel structured ncRNAs^*21, 48*^ that play important roles in both viruses, as well as multiple highly-conserved (across mammals) structures in Xist that are likely important to the mechanism of X chromosome inactivation.^*44*^ This same approach was used to discover conserved regions of structure with *MAPT* pre-mRNA that may dictate biological function.

### Global structure of tau pre-mRNA

The tau pre-mRNA has 21 predicted structured exon-intron junctions, one predicted structured region in the 5′ UTR, and nine predicted structured regions in the 3′ UTR. Altogether, approximately 1000 bps at exon-intron junctions, 60 bps in the 5′ UTR, and 2,500 bps in the 3′ UTR have structure. Five alternatively spliced exon-intron junctions have structures, which may regulate splicing. Base pair conservation at these junctions is generally high, but in relatively few non-human primates (typically less than 10). Within the 3′ UTR, eight of the nine structured regions have base pair conservation of ∼90% or greater.

### Structures at exon-intron junctions

Structures were predicted at 19 exon-intron junctions, including five of the six that are adjacent to alternatively spliced exons (Table 1 and Figures 2, 3, and S1). Base pair conservation for all of these structures was above 90%. Thus, they are highly conserved. Regulation of splicing at these exon-intron junctions may be achieved by recognition of structures or splicing elements by splicing factors. Thus, mutations within sequences near the junctions that affect structure may disrupt normal splicing and cause disease.

**Table 1:** Exon-intron junctions containing predicted wild type structures.

| Junction | Nucleotides | $\Delta G_{37}^{\circ}$<br>(kcal/mol) | Average base pair<br>conservation (%) | Number of<br>species<sup>a</sup> |
| --- | --- | --- | --- | --- |
| E1-I1 | 137-150 of E1 and<br>E1+1 to E1+7 of I1 | -9.5 | 94.3 | 5 |
| I1-E2 | E2-29 to E2-1 of I1<br>and 1-3 of E2 | -9.0 | 97.4 | 6 |
| E2-I2 | 82-87 of E2 and E2+1<br>to E2+27 of I2 | -11.6 | 99.2 | 6 |
| E2-I2 | 68-87 of E2 and E2+1<br>to E2+46 of I2 | -19.2 | 99.0 | 6 |
| E3-I3 | 18-87 of E3 | -25.9 | 92.9 | 18 |
| I3-E4 | E4-43 to E4-1 of I3<br>and 1-15 of E4 | -14.0 | 100.0 | 2 |
| I3-E4 | E4-25 to E4-1 of I3<br>and 1-6 of E4 | -6.4 | 100.0 | 1 |
| I4-E4A | E4A-38 to E4A-1 of I4<br>and 1-31 of E4A | -24.1 | 97.5 | 3 |
| I4A-E5 | E5-63 to E5-1 of I4A<br>and 1-5 of E5 | -15.4 | 100.0 | 5 |
| E5-I5 | 4-56 of E5 and E5+1 to<br>E5+15 of I5 | -13.5 | 100.0 | 2 |
| E5-I5 | 50-56 of E5 and E5+1<br>to E5+53 of I5 | -16.5 | 100.0 | 2 |
| E6-I6 | 133-198 of E6 and<br>E6+1 of I6 | -18.2 | 98.8 | 8 |
| E6-I6 | 164-198 of E6 and<br>E6+1 to E6+29 of I6 | -11.7 | 100.0 | 1 |
| I6-E7 | E7-35 to E7-1 of I6<br>and 1-28 of E7 | -14.5 | 99.0 | 4 |
| I6-E7 | E7-1 of I6 and 1-48 of<br>E7 | -17.0 | 96.3 | 9 |
| E7-I7 | E7+1 to E7+21 of I7 | -2.7 | 100.0 | 2 |
| I7-E8 | E8-19 to E8-1 of I7<br>and 1-27 of E8 | -13.0 | 100.0 | 1 |
| I7-E8 | E8-26 to E8-1 of I7<br>and 1-40 of E8 | -17.9 | 98.7 | 4 |
| I8-E9 | E9-41 to E9-1 of I8<br>and 1-21 of E9 | -11.0 | 100.0 | 4 |
| I8-E9 | E9-15 to E9-1 of I8<br>and 1-5 of E9 | -0.4 | 100.0 | 1 |
| I9-E10 | E10-14 to E10-1 of I9<br>and 1-7 of E10 | -4.5 | 94.1 | 7 |

<sup>a</sup> Non-human primates

**Table 1 (continued):** Exon-intron junctions containing predicted wild type structures.
| Junction | Nucleotides | $\Delta G_{37^\circ}$<br>(kcal/mol) | Average base pair<br>conservation (%) | Number of<br>species<sup>a</sup> |
| --- | --- | --- | --- | --- |
| E10-I10 | 73-93 of E10 and<br>E10+1 to E10+31 of I10 | -19.3 | 93.3 | 4 |
| I10-E11 | E11-3 to E11-1 of I10<br>and 1-30 of E11 | -6.4 | 94.4 | 5 |
| I10-E11 | E11-10 to E11-1 of I10<br>and 1-40 of E11 | -10.9 | 90.0 | 4 |
| E11-I11 | 78-82 of E11 and<br>E11+1 to E11+60 of I11 | -17.7 | 92.1 | 5 |
| I11-E12 | E12-61 to E12-1 of I11<br>and 1-8 of E12 | -20.1 | 94.6 | 6 |
| I11-E12 | E12-11 to E12-1 of I11<br>and 1-51 of E12 | -16.0 | 95.4 | 5 |
| E12-I12 | 77-113 of E12 and<br>E12+1 to E12+22 of I12 | -17.9 | 96.1 | 5 |
| E12-I12 | 100-113 of E12 and<br>E12+1 to E12+53 of I12 | -11.9 | 94.9 | 5 |
| I12-E13 | E13-9 to E13-1 of I12<br>and 1-32 of E13 | -6.8 | 94.0 | 7 |

<sup>a</sup> Non-human primates

**Figure 2:**
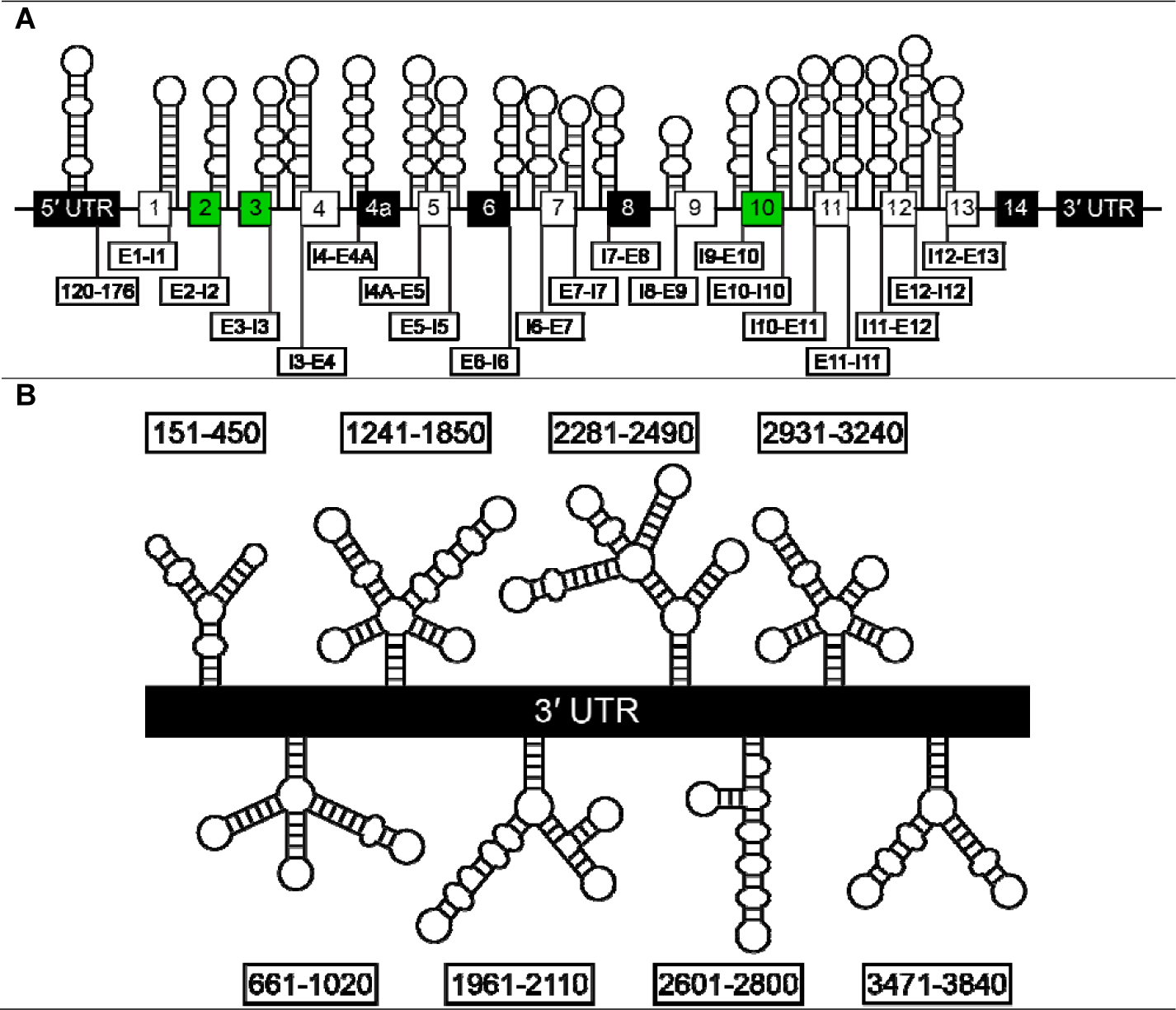
Overview of structured regions in the tau pre-mRNA. (a) Conserved structures in the 5′ UTR and exon-intron junctions. White boxes indicate constitutive exons. Black boxes indicate exons that are excluded in the human brain. Green boxes indicate exons that are alternatively spliced. (b) Conserved structures identified in the 3′ UTR.

**Figure 3:**
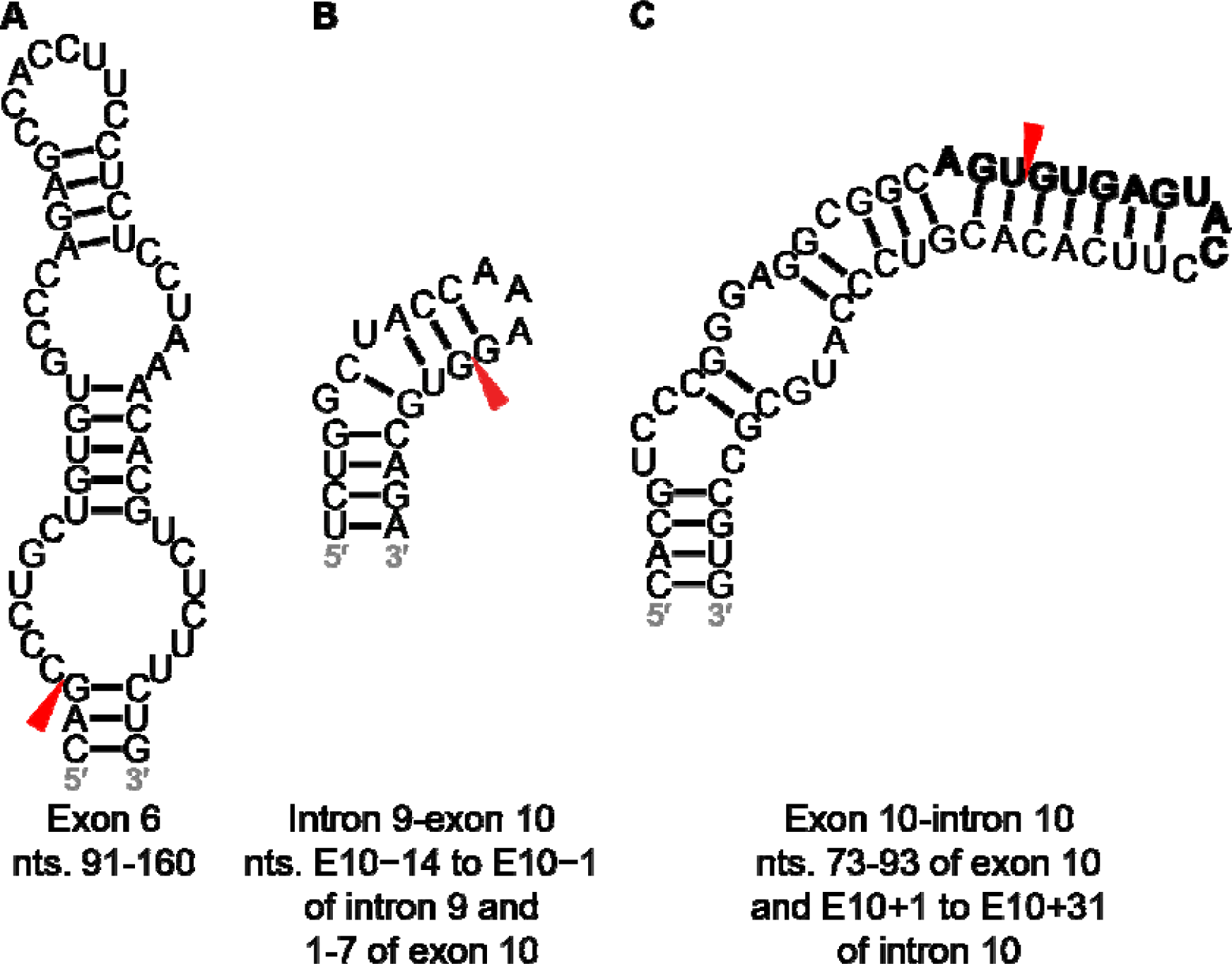
Predicted structures at alternatively-spliced exon-intron junctions. Splice sites are denoted with red arrowheads. Potential U1 snRNA binding sites are denoted by bold nucleotides. The enhancer in the hairpin structure at the exon 3-intron 3 junction is colored orange. Usage note: at a 3′ splice site, intronic nucleotides are denoted by a minus (-) sign and their position upstream of the adjacent exon, while exonic nucleotides are denoted by their position relative to the first nucleotide of the exon. At a 5′ splice site, intronic nucleotides are denoted by a (+) sign and their position downstream of the adjacent exon.

Below, we focus on the stable predicted structures at exon-intron junctions that are adjacent to alternatively spliced exons (exons 2, 3, and 10). Each contains a U1 snRNA binding site, where U1 5′ splice site recognition occurs by forming a duplex beginning at +1G of the 5′ splice site and C8 of U1 snRNA.^*49*^ A maximum of 11 bps may form between the 5′ splice site and U1 snRNA, spanning the last 3 nt of the exon and first 8 nt of the intron.^*49*^ RNA structure at the 5′ splice site may regulate splicing or compete with U1 snRNA for base pairing.^*49*^

The remainder of the structures and their associated metadata can be found in Supplementary Results and Figures S1 and S2. There is interesting biology at these other intron-exon junctions. For example, a structured region was predicted at the exon 6-intron 6 junction. Deletion mutagenesis on exon 6 revealed that the 5′ and 3′ ends, the latter of which includes a 25 – 30 nt sequence within the predicted hairpin structure, decreased exon 6 inclusion, indicating that these regions act as strong splicing enhancers.^*50*^ Exon 6 has two cryptic 3′ splice sites, which, when used, result in frameshift mutations and yield variants of tau protein lacking in microtubule binding domains.^*50, 51*^ These splice sites, termed 6p and 6d, are 101 and 170 nt long, respectively, are downstream of the canonical splice site. A stable structure was predicted at the 6p splice site (Figure 3A).^*52*^ This same structure was predicted to form from the 250 nt sequence centered on the exon 6-intron 6 junction, but its z-score and ED were within 1 SD of average for all windows.

*Exon 10.* Exon 10 alternative splicing is the most well studied splicing event in tau pre-mRNA, owing to identified mutations that cause disease (Figures 4A to 4C and 4F to 4I). ^*13, 53, 54*^ Among 70 nt windows spanning the intron 9-exon 10 junction, the window with the lowest z-score (−1.05) contains a hairpin structure modeled using SHAPE mapping (Figure 3B).^*55*^ A G-to-U mutation at position E10-10 near this junction was observed to increase the 4R-to-3R tau ratio (Figures 4D and 4E).^*56*^ The previously validated hairpin and splice site regulator at the exon 10-intron 10 junction was predicted as part of a larger hairpin spanning nt 73 – 93 of exon 10 and E10+1 – E10+31 of intron 10, which contains a U1 snRNA binding site (Figure 3C). The 70 nt window containing this structure has a z-score and ED of −1.15 and 3.39, respectively, which were more than 1 SD below their respective averages.

**Figure 4:**
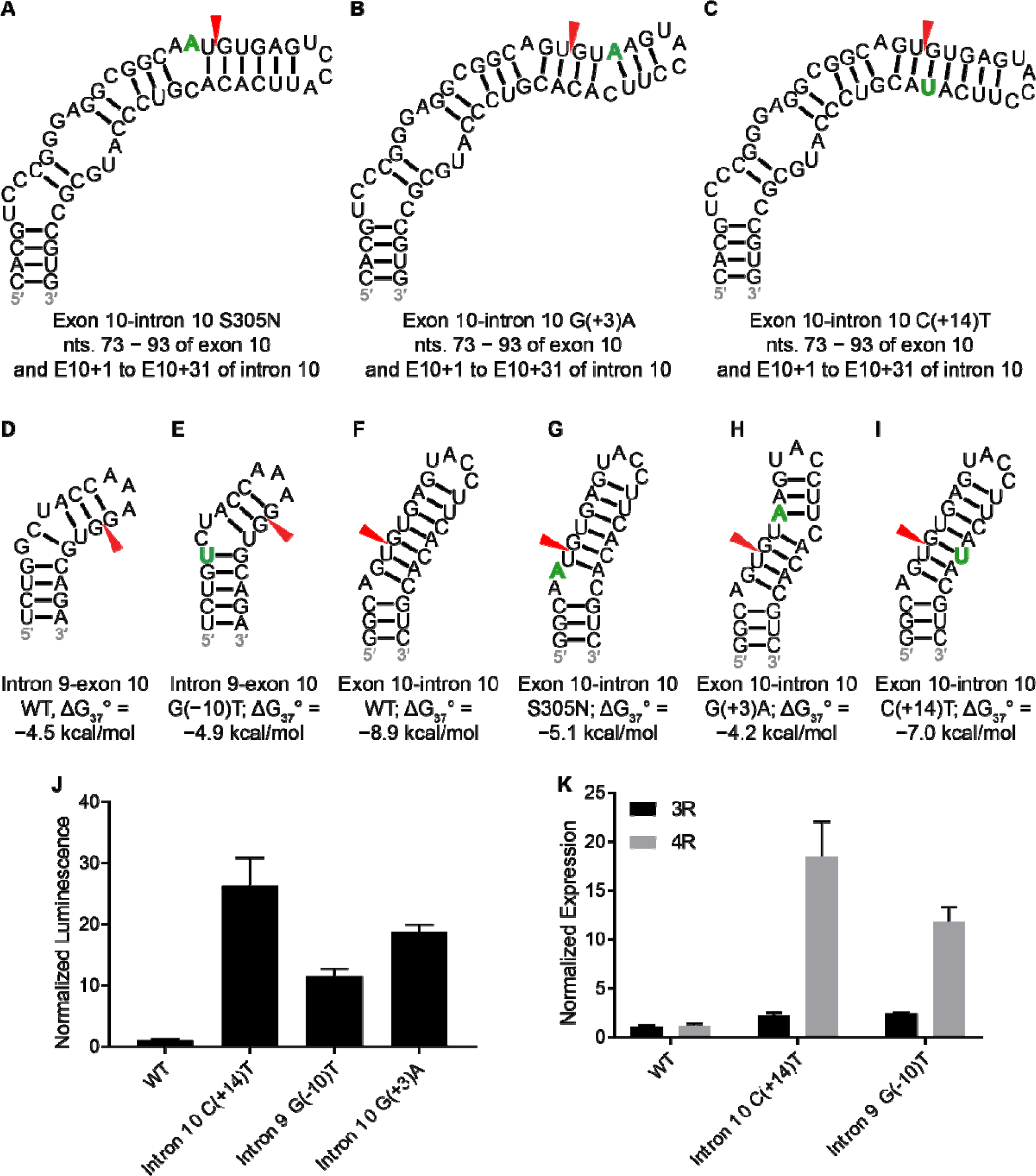
Predicted structures at exon-intron junctions that contain mutations and experimental results with minigenes at exon-intron junctions. Shown are structures containing mutations that affect the wild type structure. Mutations are denoted in green. Splice sites are denoted with red arrowheads. (a) to (c) Extended structures of hairpins at the exon 10-intron 10 junction containing S305N, G(+3)A, and C(+14)T mutations, respectively. (d) and (e) Wild type and G(−10)T mutant hairpin structures, respectively, at the intron 9-exon 10 junction. (f) to (i) Minimal wild type and mutant S305N, G(+3)A, and C(+14)T hairpin structures, respectively, at the exon 10-intron 10 junction. (j) Effect of mutations at the 5′ and 3′ splice sites of exon 10 on luciferase activity in transfected HeLa cells. (k) Effect of mutations at the 5′ and 3′ splice sites of exon 10 on 3R and 4R tau mRNA levels in HeLa cells as measured by RT-qPCR.

This hairpin structure was also supported by SHAPE mapping.^*55*^ Splicing of exon 10 is regulated by *cis*-elements, such as exonic splicing enhancers, silencers (ESEs and ESSs, respectively) and structured RNAs, and *trans*-factors, such as serine-arginine (SR) proteins that bind to ESEs adjacent to the 5′ and 3′ splice sites and promote association with U1 and U2 snRNP, respectively.^*51, 55, 57, 58*^ These splice sites are weak, and their activities (and exon 10 inclusion) are dependent on the ESEs.^*58*^ Consistent with ESE-dependent splice sites is the presence of a weak polypyrimidine tract at the 3′ splice site.^*58*^ Deletion of nt 7 – 15 of exon 10 inhibited binding and inhibition of splicing by SRp30c and SRp55.^*59*^ This region is upstream of an enhancer, defined by a 5 ′-AAG motif spanning nt 16 – 18 of exon 10 that binds splicing activator htra2β1.^*59-61*^ Deletion of this region, also known as the Δ280K mutation, decreased exon 10 inclusion and formation of 4R tau.^*59-61*^ Co-transfection of HEK293 cells with SRp54 (signal recognition particle 54) and a mutant mini-gene without nt 16 – 21 of exon 10, which consists of a purine-rich 5 ′-AAGAAG enhancer element, eliminated the binding of SRp54.^*62*^ These results indicate that htra2β1 and SRp54, which antagonizes its activity, bind to these regions.^*59-62*^

On the other hand, the N279K mutation, which changes a U to a G, increases exon 10 inclusion by strengthening an AG-rich region that acts as a splicing enhancer.^*59-61*^ Similar to exon 2, the default splicing pattern is inclusion and regulation primarily occurs through inhibition.^*51, 59*^ The S305N mutation, caused by a NG_007398.1:c.914G>A substitution (cDNA numbering starting from A in the AUG initiation codon in the noted GenBank cDNA reference sequence),^*63*^ converts a GC pair to a noncanonical AC pair (Table 2).^*64*^ The result is destabilization of the exon 10-intron 10 SRE and increased exon 10 inclusion, similar to other intronic mutations at this junction (Table 2).^*64*^ RNA2DMut predicted consistent MFE and centroid structures for this mutation, which increased ED from 3.39 to 11.16 and destabilized the structure by 3.8 kcal/mol. A G(+3)A mutation at the exon 10-intron 10 junction decreases the stability of the stem-loop by 4.7 kcal/mol by disrupting a GC pair and stabilizes formation of the U1 snRNP complex by changing a GU pair to an AU pair.^*65*^ For this mutation, RNA2DMut predicted an increase in ED of 8.02 relative to WT for the centroid structure. A C(+14)T mutation converts a GC pair in the stem-loop to a GU pair, destabilizing the helix by 1.9 kcal/mol. RNA2DMut predicted a small 0.24 increase in ED.

**Table 2:**
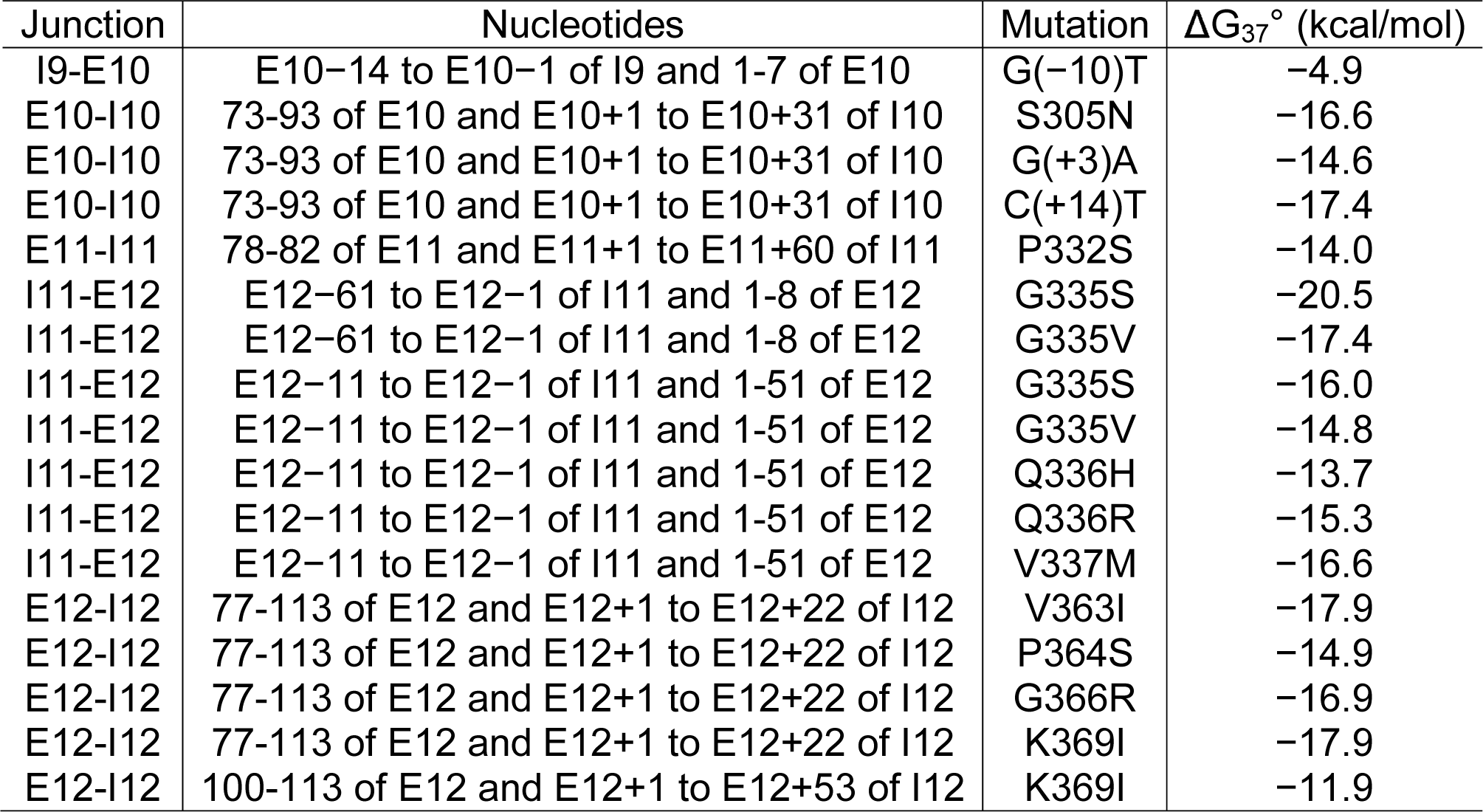
Exon-intron junctions containing predicted mutant structures.

| Junction | Nucleotides | Mutation | $\Delta G_{37}^{\circ}$ (kcal/mol) |
| --- | --- | --- | --- |
| I9-E10 | E10-14 to E10-1 of I9 and 1-7 of E10 | G(-10)T | -4.9 |
| E10-I10 | 73-93 of E10 and E10+1 to E10+31 of I10 | S305N | -16.6 |
| E10-I10 | 73-93 of E10 and E10+1 to E10+31 of I10 | G(+3)A | -14.6 |
| E10-I10 | 73-93 of E10 and E10+1 to E10+31 of I10 | C(+14)T | -17.4 |
| E11-I11 | 78-82 of E11 and E11+1 to E11+60 of I11 | P332S | -14.0 |
| I11-E12 | E12-61 to E12-1 of I11 and 1-8 of E12 | G335S | -20.5 |
| I11-E12 | E12-61 to E12-1 of I11 and 1-8 of E12 | G335V | -17.4 |
| I11-E12 | E12-11 to E12-1 of I11 and 1-51 of E12 | G335S | -16.0 |
| I11-E12 | E12-11 to E12-1 of I11 and 1-51 of E12 | G335V | -14.8 |
| I11-E12 | E12-11 to E12-1 of I11 and 1-51 of E12 | Q336H | -13.7 |
| I11-E12 | E12-11 to E12-1 of I11 and 1-51 of E12 | Q336R | -15.3 |
| I11-E12 | E12-11 to E12-1 of I11 and 1-51 of E12 | V337M | -16.6 |
| E12-I12 | 77-113 of E12 and E12+1 to E12+22 of I12 | V363I | -17.9 |
| E12-I12 | 77-113 of E12 and E12+1 to E12+22 of I12 | P364S | -14.9 |
| E12-I12 | 77-113 of E12 and E12+1 to E12+22 of I12 | G366R | -16.9 |
| E12-I12 | 77-113 of E12 and E12+1 to E12+22 of I12 | K369I | -17.9 |
| E12-I12 | 100-113 of E12 and E12+1 to E12+53 of I12 | K369I | -11.9 |

### Experimentally-determined structures at exon 10-intron junctions

The splicing regulator at the exon 10-intron 10 junction was predicted to form, with 93.3% base pair conservation (Figures 3 and 4). The C(+14)U mutation causes the disinhibition-dementia-parkinsonism-amyotrophy (DDPAC) disease, one type of FTDP-17.^*13, 53, 66*^ A 4R-to-3R ratio of 30:1 relative to wild type was measured by RT-PCR in HeLa cells transfected with mini-genes expressing the stem-loops.^*30*^ In transfected luciferase assays, the 4R-to-3R ratio of DDPAC tau to wild type tau was similar.

A C-bulge was predicted to form in the structure containing the G(+3)A mutation. This mutation results in the multiple system tauopathy with presenile dementia (MSTD) form of FTDP-17.^*54, 65*^ A 4R-to-3R ratio of ∼20:1 relative to wild type was measured in transfected luciferase assays with this mutation. No occurrence of this mutation was found in non-human primates.

A hairpin at the intron 9-exon 10 junction was predicted to form by SHAPE-constrained RNA folding algorithms (Figure 4D).^*55*^ A G-to-U mutation at position E10-10 of this splice junction (Figure 4E) weakens a polypyridine tract and a 2 nt bulge loop was predicted to form as a result of the G-to-U mutation, which stabilizes the wild type hairpin by 0.4 kcal/mol. This mutation leads to increased selection of the splice site and exon 10 inclusion.^*56, 58*^ In cell-based assays completed herein, the G-to-U mutation led to an 11.5-fold increase in luciferase expression over wild type (Figure 4J).

### Structure of the 5′ UTR

The tau 5′ UTR contains a conserved oligopyrimidine sequence, 5′-CCTCCCC-3′ that promotes cap-dependent translation by activation of the mTOR pathway.^*67, 68*^ Separately, eukaryotic internal ribosome entry sites (IRESs) allow initiation of cap-independent translation of mRNA.^*69, 70*^ IRES activity may be mediated by RNA structure, short sequence motifs in unstructured or structured regions that bind to IRES trans-acting factors (ITAFs), or base pairing with 18S ribosomal RNA (rRNA).^*69, 70*^ Additionally, sequence motifs within loop regions of IRESs may allow for RNA-RNA interactions and formation of secondary or tertiary structure essential for IRES function.^*69, 70*^ The tau 5′ UTR contains a cis-regulatory internal ribosomal entry site (IRES) that contributed to 30% of the total translation of tau mRNA in luciferase assays.^*67-69*^ Truncations to the IRES reduced or abolished IRES activity, suggesting that the entire sequence is required for IRES activity.^*69*^ A secondary structure model of the 240 nt IRES generated by SHAPE analysis yielded two domains. Among them, Domain I is a hairpin and Domain II is a multibranch loop.^*69*^ Mutations that disrupted the stem regions of Domains I and II abolished IRES activity.^*69*^ Therefore, sequences and/or structures in these regions are required for IRES function.^*69*^

Our analysis predicts a structured region corresponding to nt 120 – 176 of the 304 nt 5′ UTR, which lies within Domain I of the IRES and has a favorable z-score and low ED (−1.03 and 5.82, respectively) (Figure 5). The structure is similar to Domain I, although the base pairs around the internal loop below the apical loop were shifted, resulting in a 3 × 1 instead of 3 × 4 internal loop. This may result from the lone cytidine on the 3′ side of the loop having SHAPE reactivity in experiments with transfected SK-N-SH cells, whereas other neighboring nucleotides were unreactive.^*69*^ The free energies of 42 nt segments of the unconstrained and SHAPE-constrained Domain I hairpin structures (−21.0 and −20.6 kcal/mol, respectively) are relatively close. Therefore, these conformations may exist in equilibrium. The probabilities of formation of loops in the predicted and SHAPE-constrained 5′ UTR was analyzed with ProbScan as a measure of the accuracy of structure prediction compared to experiment.^*27*^ The probabilities of the 3 × 4 internal loop and 9 nt hairpin loop calculated by ProbScan for the predicted structure are 0.584 and 0.601, respectively. Folding of the same fragment of sequence of the predicted structure into the SHAPE-constrained 3 × 1 internal loop and 10 nt hairpin loop resulted in loop probabilities of 0.301 and 0.323, respectively. These results suggest that the predicted 3 × 4 loop and associated hairpin loop may be more likely to form than those in the SHAPE-constrained structure and are consistent with the slightly more favorable folding free energy of the conformation of the predicted structure relative to the SHAPE-constrained structure. When the full 5′ UTR was queried, a region within the human structure corresponding to base pairs C133:G160 to G142:C150 and containing a 3 × 4 internal loop was predicted from alignment and folding of the resulting sequences. Base pair conservation for this consensus structure is 94.6% in up to 20 non-human primates. Altogether, a hairpin structure consistent with a SHAPE-constrained IRES structure was predicted to form in the 5′ UTR and contains structural and sequence elements required for IRES activity.

**Figure 5:**
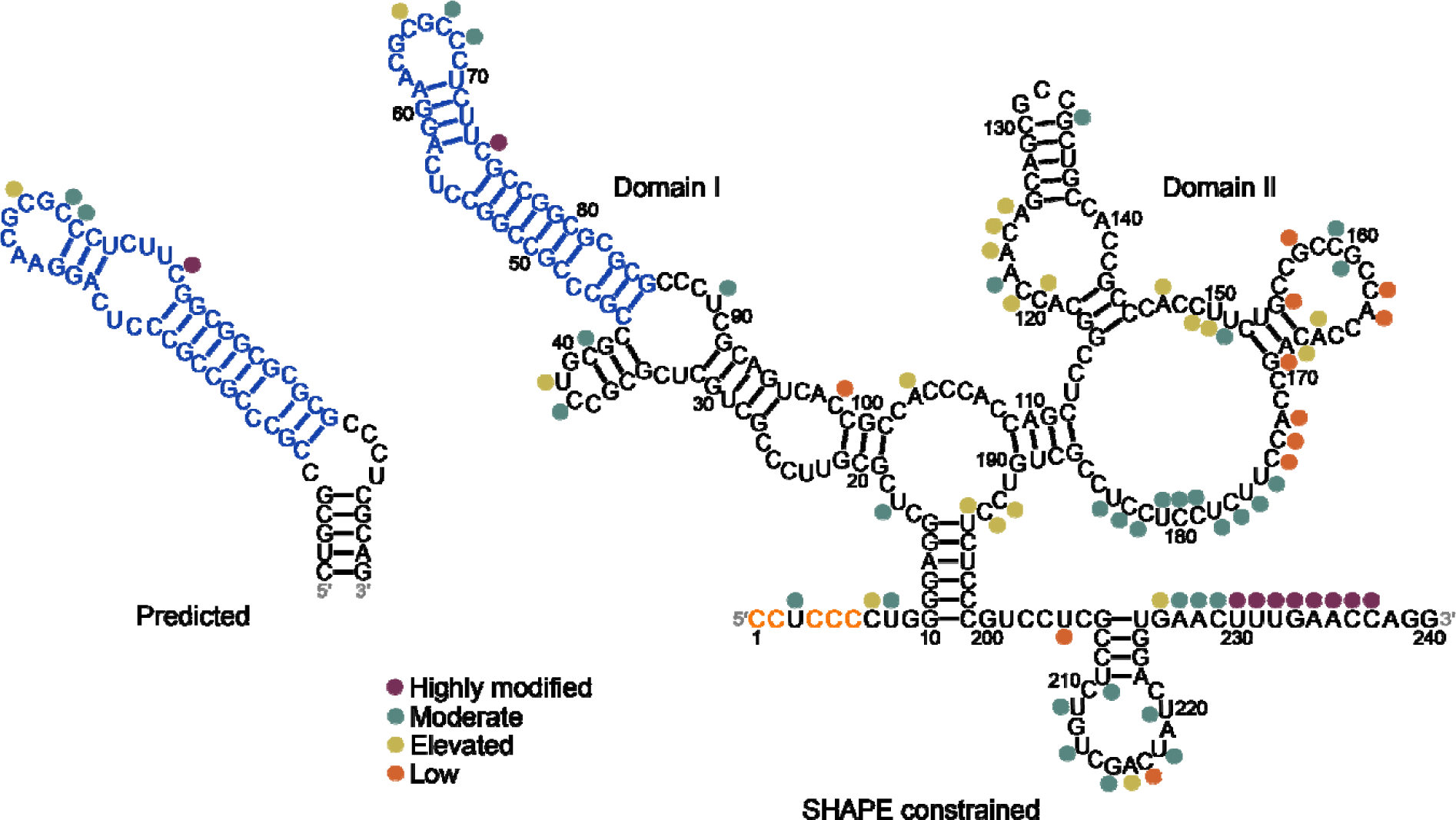
Predicted and SHAPE constrained structures of the human 5′ UTR. Predicted structure corresponds to nt 120 – 176. Structure determined by SHAPE analysis corresponds to the 240 nt tau IRES.^*69*^ The conserved 5′-CCUCCCC-3′ sequence that promotes cap-dependent translation is colored orange. Blue nucleotides in the SHAPE constrained structure correspond to those in the predicted structure. Nucleotides in the SHAPE constrained structure that were modified in SHAPE analysis are indicated by circles colored according to SHAPE reactivity (see figure key). For reference, nucleotides in the predicted structure that correspond to those in the SHAPE-constrained structure are labeled with the respective colored circles. SHAPE reactivities were obtained from ref. *69*.

### Structure of the 3′ UTR

The 3′ UTR of mRNAs regulate various cellular events such as translation, and transcript stability, polyadenylation, and localization.^*71*^ One mechanism by which regulation of gene expression occurs is by alternative polyadenylation (APA).^*22, 28, 72, 73*^ Polyadenylation at alternative sites in the 3′ UTR results in transcripts with 3′ UTRs of different lengths.^*72*^ Longer 3′ UTRs may contain more miRNA binding sites or regulatory sequences that cause the mRNA to be more prone to negative regulation.^*72*^ The human tau 3′ UTR contains two polyadenylation signals, which results in 3′ UTR isoforms 256 nt and 4163 nt in length.^*15*^ Dickson *et al.* found that the short isoform was expressed significantly higher than the long isoform.^*15*^ The differential regulation of the 3′ UTR isoforms may be mediated by *cis*-elements and *trans*-factors.^*15*^ Dickson *et al.* identified a binding site for miR-34a, which inhibits expression of endogenous tau, and reported that a luciferase reporter vector containing a mutant miR-34a binding site was insensitive to pre-miR-34a.^*15*^ Additionally, cloning of fragments of the long 3′ UTR isoform into a luciferase reporter vector increased or decreased luciferase activity in transfected M17D and SH-SY5Y cells, suggesting that these fragments may differentially affect tau expression.^*15, 74*^ Secondary structures within the 3′ UTR are vital determinants of gene levels, and mutations affecting secondary structure or a polyadenylation signal can cause disease.^*71*^

Indeed, the tau 3′ UTR has been implicated in stabilization of tau mRNA in neuronal cells.^*75*^ The ratio of 3′ UTR to coding DNA sequence is balanced in normal brain tissue but elevated in the AD brain, suggesting that the 3′ UTR may have regulatory roles in the disease.^*74*^ The 3′ UTR contains several microRNA binding sites such as miR-34a, miR-132, miR-181c, miR-219, miR-485-5p, and miR-642 (Table S2).^*15, 74, 76, 77*^ In particular, expression of miR-219-5p and miR-485-5p are decreased in AD and may be attributed to overexpression of tau.^*74, 76-78*^ Thus, targeting structured regions containing microRNA binding sites may be an approach for treating tauopathies.

Eight regions within the long 3′ UTR isoform were predicted to have z-scores less than 1 SD below the average of all windows (Table 3). Six of the regions had base pair conservation above 90%. Positions 151 – 300 have an average canonical base pair conservation of 97.6%. Within this region is a 5′-AAUAAA polyadenylation signal, which lies within a 5 × 5 internal loop and spans nt 230 – 235 of the 3′ UTR.^*31*^ Cleavage and polyadenylation at this site generates the short 3′ UTR isoform.^*31*^

**Table 3:** Regions of the 3′ UTR with predicted structure. Secondary structures for these regions are in Figures 6 and S3 – S8.

| Nucleotides | $\Delta G_{37^\circ}$<br>(kcal/mol) | Average base<br>pair conservation<br>(%) |
| --- | --- | --- |
| 151-450 | -91.1 | 97.7 |
| 661-1020 | -147.1 | 94.0 |
| 1241-1850 | -219.6 | 93.8 |
| 1961-2110 | -50.2 | 89.2 |
| 2281-2490 | -81.7 | 95.2 |
| 2601-2800 | -66.0 | 43.7 |
| 2931-3240 | -98.3 | 89.2 |
| 3471-3840 | -149.6 | 94.9 |

The structured region predicted to form between nt 1961 – 2110 in the 3′ UTR (average canonical base pair conservation of 89.2%) is similar to the multibranch loop formed in the tau coding region (nt 1989-2015). Positions 2281 – 2490 have an average canonical base pair conservation of 95.2%. These results suggest that this region is highly likely to form functional structure.

Other regions in the 3′ UTR are also highly conserved, which span (or are adjacent to) microRNA binding sites, nt 661 – 1020 (94.0%) and nt 1241 – 1850 (93.8%). Interestingly, deletion of nt 986 – 1479, which includes binding sites for miR-34a and miR-485-5p and partly overlaps with this region, from a luciferase reporter construct significantly increased luciferase activity in transfected HEK293 and differentiated SH-SY5Y cells compared to cells expressing a wild type 3′ UTR construct.^*31*^ This suggests that this region may contain a *cis*-element that downregulates expression of tau.^*31*^ Indeed, a binding site for miR-485-5p is located upstream of the structured region comprised by nt 1241 – 1850, in addition to the RNA structure itself.^*76*^ Together, these may control expression of tau.

Two structured regions, corresponding to nt 1961 – 2110 and 3471 – 3870, lie in segments that upregulated luciferase activity in transfected cells reported by Dickson et al (Figures 6A and B).^*15*^ In cell-based assays carried out in this work, three fragments of the 3′ UTR were found to upregulate luciferase activity (Figure 6C). These three fragments, corresponding to nt 2041 – 2079, 3471 – 3691, and 3695 – 3859, contain regions that were predicted to be structured. In other studies, deletion of nt 3501 – 4000 increased luciferase activity in cells relative to those expressing WT tau.^*31*^ Therefore, the predicted structures may contain binding sites for *cis*-elements or trans factors that regulate tau expression.

**Figure 6:**
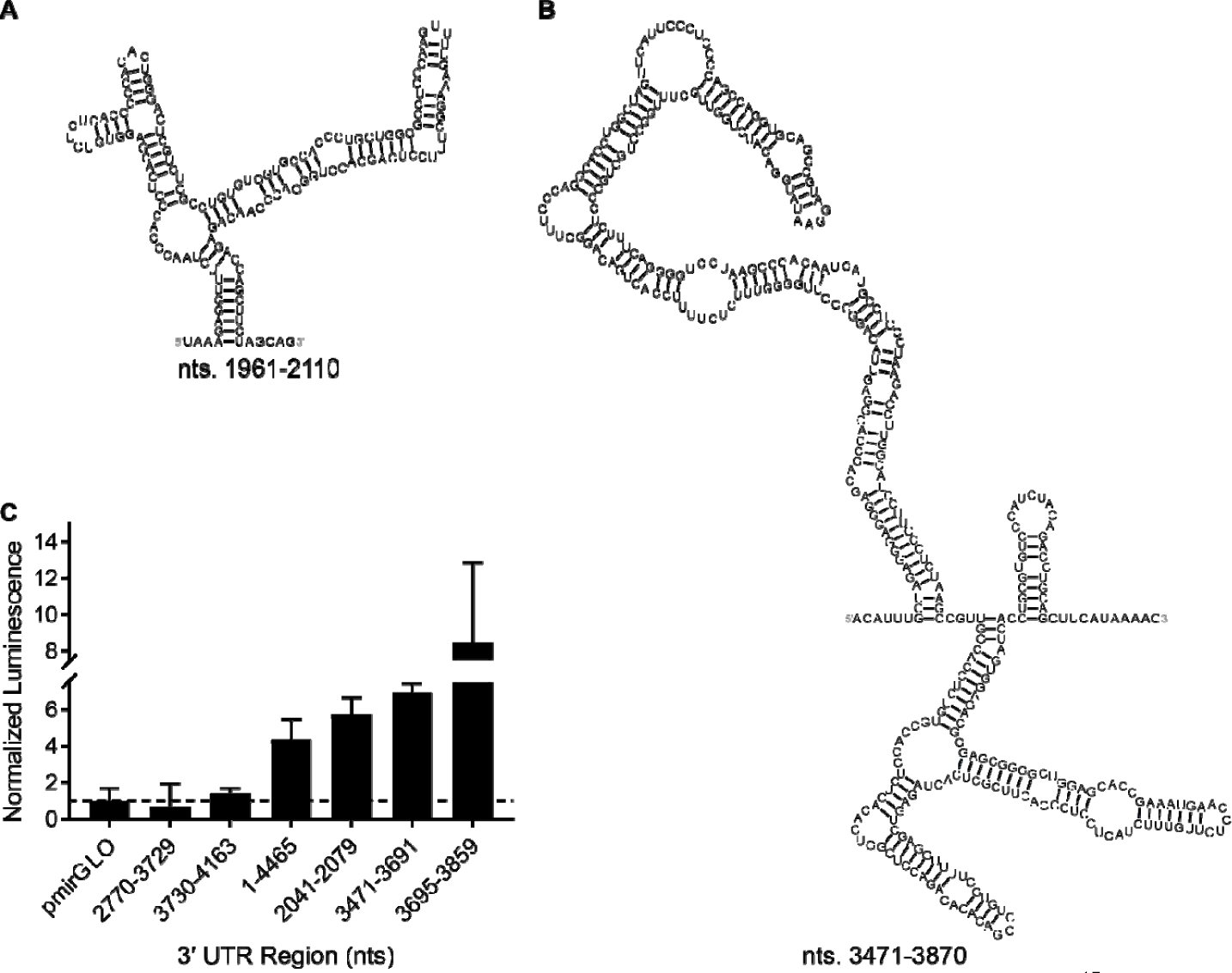
Structured regions of the 3′ UTR that upregulated luciferase activity in Dickson *et al*^*15*^ and results of cell-based luciferase assays completed herein with fragments of the 3′ UTR.

## CONCLUSIONS

In summary, stable structures were predicted throughout the *MAPT* pre-mRNA. Structures were predicted at 19 exon-intron junctions, including five adjacent to alternatively spliced exons (Figure 1). Two folds were predicted at the exon 2-intron 2 junction, the larger of which is an extended form of a hairpin where the stem contains two single nucleotide bulges. The predicted structures at the 5′ and 3′ splice sites of exon 10 were previously identified by experiment. The G(+3)A and C(+14)T mutations that are associated with disease were verified by cell-based assays to upregulate exon 10 inclusion. These mutations destabilize the hairpin structure at the exon 10-intron 10 junction. A structure was predicted to form in exon 6 at a cryptic splice site, which results in frameshift. The IRES within the 5′ UTR contains a predicted structure, mutations within which had been demonstrated to decrease IRES activity. Eight structured regions were identified throughout the 3′ UTR, including regions that regulate expression of tau. RNA secondary structure plays important roles in regulating gene expression, and splice site junctions and UTRs are known to contain gene regulatory element. These regions of pre-mRNA may interact with *cis*-elements or *trans*-factors via sequence- or structure-specific binding. The results of this study may be applied to future studies to identify new therapeutic targets against tau-associated diseases.^*32, 79-82*^

## ABBREVIATIONS

AD: Alzheimer’s disease
APA: alternative polyadenylation
APSI: average pairwise sequence identity
bp: base pair
DMEM: Dulbecco’s Modified Eagle Medium
ED: ensemble diversity
FTDP-17: frontotemporal dementia with parkinsonism-17
htra2β1: transformer 2 beta homolog 1
IRES: internal ribosome entry site
ITAF: IRES trans-acting factor
MAFFT: Multiple Alignment using Fast Fourier Transform
MAPT: microtubule associated protein tau
MBD: microtubule binding domain
MFE: minimum free energy
miRNA: microRNA
mRNA: messenger RNA
mTOR: mammalian target of rapamycin
MSTD: multiple system tauopathy with presenile dementia
ncRNA: noncoding RNA
nt: nucleotide
PD: Parkinson’s disease
RNP: ribonucleoprotein
rRNA: ribosomal RNA
SCI: structure conservation index
SD: standard deviation
SF2: serine and arginine rich splicing factor 1 also known as SRSF1
SHAPE: selective 2′ hydroxyl acylation analyzed by primer extension
snRNA: small nuclear RNA
SRp30c: serine and arginine rich splicing factor 9 also known as SRSF9
SRp40: serine and arginine rich splicing factor 5 also known as SRSF5
SRp54: signal recognition particle 54
SRp55: serine and arginine rich splicing factor 6 also known as SRSF6
SVM: support vector machine
UTR: untranslated region

## ACKNOWLEDGEMENTS

This work was supported by the National Institutes of Health (R01-GM097455-07, DP1-NS096898, and P01-NS099114 to M.D.D and R00-GM112877 to W.N.M) and the Tau Consortium and Rainwater Charitable Foundation (to M.D.D.) as well as startup funds from the Iowa State University College of Agriculture and Life Sciences and the Roy J. Carver Charitable Trust (W.N.M.) and the Huntington’s Disease Society of America (J.L.C.).

