## Supplementary Information for "The RNA encoding the microtubule-associated protein tau has extensive structure that affects its biology"

Jonathan L. Chen,^1^ Walter N. Moss,^2^ Adam Spencer,^1^ Peiyuan Zhang,^1^ Jessica L. Childs-Disney^1^ and Matthew D. Disney^1,^*

^1^ Department of Chemistry, The Scripps Research Institute, Jupiter, FL 33458, United States.

^2^ Roy J. Carver Department of Biochemistry, Biophysics & Molecular Biology, Iowa State University, Ames, IA 50011, United States.

Table S1	Primers used to clone fragments of tau 3′ UTR into a pmirGLO vector.

Table S2	MicroRNAs predicted to bind to the tau 3′ UTR.

Figure S1	Predicted structures at other exon-intron junctions.

Figure S2	Predicted structures at exon-intron junctions that contain mutations.

Figure S3	Positions 151-450 of the 3′ UTR.

Figure S4	Positions 661-1020 of the 3′ UTR.

Figure S5	Positions 1241-1850 of the 3′ UTR.

Figure S6	Positions 2281-2490 of the 3′ UTR.

Figure S7	Positions 2601-2800 of the 3′ UTR.

Figure S8	Positions 2931-3240 of the 3′ UTR.

**SUPPLEMENTARY METHODS**

**Cloning of fragments of tau 3′ UTR into a pmirGLO vector.** DNA oligonucleotides with sequences corresponding to the fragment nt 2041-2077 of the 3′ UTR was purchased from Integrated DNA Technologies (IDT). PCR primers for fragments 3471-3691 and 3695-3859 were also purchased from (IDT). Oligonucleotides for 2041-2077 were annealed by heating to 95 °C for 10 mins and then cooled to room temperature, and then purified using a DNA cleanup kit. The annealed oligonucleotide was phosphorylated at the 5′ termini using T4 polynucleotide kinase. Primers corresponding to fragments nt 3471-3691 and 3695-3859 were used to PCR amplify the fragment from full length UTR (1X Q5 Reaction Buffer (NEB) using 500 nM forward primer, 500 nM reverse primer, 2 μl 25X dNTP mix, and 0.5 μl Q5 High-Fidelity DNA polymerase, in a 50 μl reaction (see Table S1 for primer sequences). PCR cycling conditions were initial denaturation at 98 °C for 30 s, then 34 cycles of 98 °C for 30 s, 60 °C for 30 s, and 72 °C for 20 s, then final extension at 72 °C for 2 mins, and then hold at 4 °C. Following cleanup of PCR products, PCR products and the pmirGLO plasmid containing the insert for 3′ UTR nts 2770-3729 (from Prof. Michael S. Wolfe) were cut using XhoI and SalI-HF in CutSmart Buffer (NEB) at 37 °C for 2 hrs and then gel purified. Inserts were ligated into the pmirGLO vector at 1:3 vector:insert ratio in a 20 μl reaction for 10 mins at room temperature. Ligated plasmids were introduced into *E. coli* DH5α cells for replication.

**SUPPLEMENTARY RESULTS**

***Structures at exon-intron junctions***

A hairpin with a fully base paired stem was predicted at the exon 1-intron 1 junction (Figure S1A). The z-score of −2.13 for the 70 nt window containing this hairpin was the lowest among all windows at this exon-intron junction and considerably below their respective averages. This window size was used as a compromise between larger window sizes, which may not predict small, regulatory motifs at exon-intron junctions, and smaller window sizes, which may not predict motifs present in larger, more stable structures. Base pair conservation in the consensus structure was predicted in nonhuman primates. Interestingly, the predicted hairpin contains a U1 snRNA binding site that spans the hairpin loop and 3′ side of the stem.*^1^* Thus, RNA structure here may be modulating accessibility of this region to U1 or other splicing regulatory factors.

*Exon 2.* Splicing of exon 2, which is included in most cells, is regulated by a purine-rich enhancer within the exon.*^2^**^,^* *^3^* The exon is regulated by CELF4 and suppressed in myotonic dystrophy type 1 (DM1), where CTG repeats sequester CELF family proteins.*^2^**^,^* *^3^* Indeed, a hairpin structure was predicted at the intron 1-exon 2 junction with a z-score and ED score of −0.89 and 7.01, respectively, within the 250 nt sequence centered on the exon 2-intron 2 junction, but not the sequence centered on the intron 1-exon 2 junction (Figure S1B). In cell-based assays reported in the literature, deletion of nt 46 – 52 and 76 – 83, which overlap with predicted structure at this junction, significantly decreased exon 2 inclusion, indicating that these sequences contain enhancers.*^2^* Deletion of nt 2 – 7 and 9 – 15, which overlap with predicted structure at the intron 1-exon 2 junction, increased exon 2 inclusion, indicating that the first ~20 residues of exon 2 is a pyrimidine-rich silencer.*^2^* Both of these deletions reduced binding to SRp30c (serine and arginine rich splicing factor 9, also known as SRSF9) while deletion of nt 9 – 15 also decreased binding to SRp55 (serine and arginine rich splicing factor 6, also known as SRSF6).*^2^* Furthermore, this region contains a sequence close to the consensus recognition sequence for SRp55.*^2^* Immunoprecipitation of these factors, in addition to htra2β1 (transformer 2 beta homolog 1), indicate that that SRp30c and SRp55 form a complex with htra2β1 and bind directly to the silencer at the 5′ end of exon 2 to modulate splicing.*^2^*

At the exon 2-intron 2 junction, a hairpin with two 1 nt bulge loops was predicted in a 70 nt window with a z-score and ED of −1.46 and 6.19, respectively (Figure S1C). In this structure, which consists of nt 82 – 87 of exon 2 and E2+1 to E2+27 (nt 1 – 27 of intron 2, downstream of the exon 2-intron 2 junction) of intron 2, the splice site lies in a helix between the two 1 nt bulge loops (Figure S1D). Both single nucleotide bulges and the first two nucleotides of the hairpin loop are part of a U1 snRNA binding site. This hairpin also appeared in a separate window, which contains nt 46 to 87 of exon 2 and E2+1 to E2+28 (nt 84 – 153 of the 250 nt sequence) with a z-score and ED of −1.15 and 8.38, respectively.

*Exon 3.* Inclusion of exon 3 is enhanced by splicing of exon 2.*^3^* Exon 3, by default, is excluded due to a weak branch point upstream of the 3′ splice site, which is attributed to poor agreement with the consensus sequence and short polypyrimidine tract.*^3-5^* Substitution with a strong branch point resulted in constitutive inclusion of the exon.*^3-5^* Directed mutagenesis experiments revealed two silencers and one enhancer within exon 3, including an enhancer at the 5′ splice site that binds to U1 snRNA (sequence is 5′-CAGgtgagg) and overlaps with a stem region of the predicted hairpin.*^4^* Exon 3 is never incorporated independently of exon 2, which may be explained by the preference of the 5′ splice site of exon 2 for the 3′ splice site of exon 3 in the absence of other competing splice sites.*^3-5^* The exon 3-intron 3 junction was predicted to contain a hairpin (z-score and ED of −1.13 and 6.09, respectively) with a 4 × 8 and a 9 × 6 nt internal loop, although the splice site junction was not predicted to be part of any structure (Figure S1E). The cytosine of the closing base pair on the 3′ side of the hairpin, however, is part of a U1 snRNA binding site. The consensus structure in this window is more conserved than others near exon-intron junctions.

At the intron 3-exon 4 junction, a hairpin containing 7 nt and 3 nt bulge loops was found and corresponds to nt E4−43 – E4−1 and nt 1 – 15 of exon 4 (Figures S1F and S1G). The z-score and ED of −2.43 and 5.41, respectively, for this 70 nt window are more than one SD below their respective averages. The upper portion of this hairpin, which consists of nt E4−25 – E4−1 and nt 1 – 6 of exon 4, also lies in the window with the lowest z-score for the 250 nt sequence at this exon-intron junction.

A hairpin containing a 5 × 4 and a 6 × 6 nt internal loop was predicted at the intron 4-exon 4A junction (Figure S1H). Exon 4A is not found in human tau.*^3^**^,^* *^5^* At the intron 4A-exon 5 junction, a hairpin was predicted with a z-score of −2.24 (Figure S1I). However, the predicted ED (5.9) was within 1 SD (5.9), indicating a potential for multiple conformations. Two separate structures, one containing nt 4 – 56 of exon 5 and E5+1 – E5+15 and another containing nt 50 – 56 of exon 5 and E5+1 to E5+53 (in windows nt 73 – 142 and 109 – 178, respectively), were predicted at the exon 5-intron 5 junction with similar z-scores (−1.62 and −1.66, respectively) (Figures S1J and S1K). Both internal loops in the first structure, in addition to the 5′ side of the 4 × 6 internal loop of the second structure, are part of a sequence recognized by U1 snRNA. However, the ED score of the second structure, which consists of the last 7 nt of exon 5 and first 53 nt of intron 5, is lower (3.03 vs. 8.97 for the first structure), and its free energy is slightly more favorable (−16.4 vs. −14.3 kcal/mol), which results in a factor of 30 in K_eq_; indicating that this may be the predominant conformation.

At the exon 6-intron 6 junction, two different structures with z-scores more than 1 SD below average were predicted (Figure S1L and S1M). The first structure, which consists of nt 133 – 198 of exon 6 and nt E6+1 (within the nt 58 – 127 window), contains a hairpin closed by a 3 nt loop while the second structure, which consists of nt 164 – 198 of exon 6 and nt E6+1 to E6+29 (within the nt 86 – 155 window), contains a multibranch loop. The first structure at this junction, however, had a lower ED score (3.47 vs. 6.29) and free energy (−18.9 vs. −14 kcal/mol) than the second structure, which corresponds to a factor of 2800 in K_eq_. In the hairpin structure, the U1 snRNA binding site is within the internal loop adjacent to the terminal helix. In the multibranch loop structure, the U1 snRNA binding site spans the multibranch loop and one of the hairpin loops. Deletion mutagenesis on exon 6 revealed that the 5′ and 3′ ends, the latter of which includes a 25 – 30 nt sequence within the predicted hairpin structure, decreased exon 6 inclusion, indicating that these regions act as strong enhancers.*^6^*

At the intron 6-exon 7 junction, two different hairpins were predicted with similar ED scores (9.36 and 9.43), although they were within 1 SD of the average for all 70 nt windows at this junction (Figures S1N and S1O). The first hairpin, which contains nt E7 − 35 of intron 6 and 1 – 28 of exon 7, has a z-score of −1.47 while the second hairpin, which contains nt E7−1 of intron 6 and 1 – 48 of exon 7, has a slightly more favorable z-score of −1.63 and lower free energy (−17 vs. −14.5 kcal/mol), which represents a factor of 58 in K_eq_. A stem-loop closed by a 4 nt hairpin loop and containing a 1 nt bulge loop was predicted near the exon 7-intron 7 junction (Figure S1P). A U1 snRNA binding site consists of the helix terminus and A-bulge. The z-score and ED scores for this window (−1.81 and 6.4, respectively) were more than 1 SD below their respective averages (−0.60 ± 1.17 and 13.14 ± 5.10, respectively) for all windows. This indicates a high propensity for being ordered to fold and a single dominant conformation.

Similar to the exon 2-intron 2 junction, a hairpin loop was predicted in two different windows at the intron 7-exon 8 junction with favorable z-scores (Figures S1Q and S1R). A 46 nt hairpin containing a 6 nt bulge loop was predicted in a window that spans nt E8−19 – E8−1 of intron 7 and 1 – 27 of exon 8. This window had a z-score of −2.7 and ED score of 6.09, which were more than 1 SD below their respective averages for all windows in the 250 nt sequence at this junction. An extended form of the hairpin was predicted in a window that spans the nt E8−26 – E8−1 of intron 7 and 1 – 40 of exon 8, and contains an additional 7 nt bulge loop. The z-score (−1.61) and ED (10.47) for this window, however, were higher than for the previous window, which contains an additional hairpin upstream of the splice site (not shown), and its free energy was slightly less favorable (−13.0 vs. −17.9 kcal/mol), resulting in a factor of 4900 in K_eq_. Exon 8 has not been found in human tau isoforms.*^3^*

Two hairpins were predicted in different 70 nt windows with favorable z-scores at the intron 8-exon 9 junction (Figures S1S and S1T). However, the ED scores for these windows (14.46 and 11.72) were relatively close to average for all windows of the 250 nt sequence (14.11 ± 5.85) and relatively high for structured regions. The hairpin that spans nt E9−41 – E9−1 of intron 8 and 1 – 21 of exon 9 contains a 5′-CAGGGGA enhancer sequence, spanning nt 16 – 22, within exon 9 that binds SF2 and SRp40 (serine and arginine rich splicing factor 1).*^7^* However, targeting this sequence and the 3′ splice site of exon 9 did not reduce tau levels.*^7^* Within exon 9, an I260V mutation, corresponding to an A-to-G change at position 222 in exon 9, increases 4R tau aggregation and decreases microtubule assembly.*^8^*

At the intron 10-exon 11 junction, a structure is predicted in two different 70 nt windows (Figures S1U and S1V). The first window, which corresponds to nt E11−39 – E11−1 and nt 1 – 31 of exon 11, had the lowest z-score (−1.69) and ED (1.54) among windows that spanned the splice junction. A structure containing a 3 × 3 internal loop and closed by a 6 nt hairpin loop was predicted with a free energy of −6.4 kcal/mol. The second window, which corresponds to nt E11−26 – E11−1 and nt 1 – 44 of exon 11, contained an extended form of this hairpin with an additional 1 × 4 internal loop and had a slightly less favorable z-score of −5.4. The additional structure in the hairpin in this window resulted in a more favorable free energy (−10.9 kcal/mol) compared to that in the first window, but its ED was higher (5.4) and within 1 SD of the average for all windows.

At the exon 11-intron 11 junction, a hairpin was predicted with z-score and ED −2.11 and 5.54, respectively, which were more than 1 SD below their averages for the 250 nt sequence (Figure S1W). The internal loop in this structure contains part of a U1 snRNA binding site. A P332S mutation (Figure S2A) near this splice junction, which caused by a NG_007398.1:c.994C>T substitution and reduces the capacity of tau to bind to microtubules, converts a GC pair to a GU pair.*^9^* The result is destabilization of the structure by 2.6 kcal/mol and an increase in ED of 4.61, from 5.54 to 10.15.

Stable structures were also predicted in the intron 11-exon 12 junction, the splice site of which lies in a helical region. The G335S and G335V mutations in this region, which are caused by GGC to AGC and GTC transitions, respectively, reduce the ability of tau to promote microtubule assembly.*^10^**^,^* *^11^* The G335S mutation converts a GU pair to an AU pair, and the G335V mutation destabilizes the terminal helix of the hairpin loop (Table 2). Two different hairpin structures were predicted at the intron 11-exon 12 junction (Figures S1X and S1Y). The first hairpin, which contained nt E12−61 – E12−1 and nt 1 – 8 of exon 12, had a z-score and an ED of −1.98 and 4.2, respectively, and its free energy of −20.1 kcal/mol was among the most favorable at this splice junction. The second hairpin, which contained nt E12−11 – E12−1 and nt 1 – 51 of exon 12, had a less favorable z-score (−1.44), ED (8.8), and free energy (−16.6 kcal/mol), which would be predicted to yield a 4100-fold difference in K_eq_. The splice junctions in these structures lie within helical regions. G335S and G335V mutations, which are caused by GGC to AGC and GTC transitions, respectively (Figures S2B and S2C), reduce the ability of tau to promote microtubule assembly.*^10^**^,^* *^11^* In both of the predicted structures at the intron 11-exon 12 junction, the G335S mutation converts a GU pair to an AU pair. In the structure consisting of nt E12−61 – E12−1 and nt 1 – 8 of exon 12, the G335V mutation destabilizes the terminal helix of the hairpin loop, resulting in 1.2 kcal/mol increase in ΔG° and an increase in ED of 1.59, from 4.2 to 5.79 (Table 2). In the structure containing nt E12−11 – E12−1 and nt 1 – 51 of exon 12, the mutation results in considerable structural changes above the splice site, including eliminating a 2 × 3 internal loop and introducing an A-bulge and a GA pair (Table 2). The result is destabilization of the structure by 1.2 kcal/mol and an increase in ED by 3.49 (8.8 to 12.29) for the RNA2DMut-predicted centroid structure relative to WT. Q336H and Q336R mutations (Figures S2D and S2E, respectively), which are caused by CAG to CAC and CGG transitions, respectively, at codon 336, promote microtubule assembly.*^12^**^,^* *^13^*

In the structure containing nt E12−11 – E12−1 and nt 1 – 51 of exon 12, the mutation causes structural changes above the 2 × 3 internal loop and introduces a 4 × 4 internal loop to the hairpin. The resulting structure is 2.3 kcal/mol less stable than wild type. In comparison, the Q336R mutation causes no change to the structure of the same hairpin except for converting an AU pair to a GU pair. RNA2DMut predicted a decrease in ED to 6.72 for the Q336H mutation and an increase in ED to 9.48 for the Q336R mutation. The V337M mutation, caused by a G-to-A substitution at nucleotide 1009, causes increased phosphorylation of tau, formation of neurofibrillary tangles and, consequently, FTDP-17 (Figure S2F).*^14-16^* The E342V mutation, associated with FTD, is caused by an A-to-T change in codon 342, which may disrupt splicing enhancers.*^17^* Predicted structural changes caused by the V337M and E342V mutations are limited to transitions from AU to GU pairs, or vice versa, but with a decrease in ED to 7.27 and an increase in ED to 8.85 for these respective mutations.

Two hairpins with favorable z-scores and low ED were predicted at the exon 12-intron 12 junction (Figures S1Z and S1AA). The window containing the first structure, which contains nt 77 – 113 of exon 12 and E12+1 to E12+22 of intron 12, has a free energy around 6.0 kcal/mol more favorable than the second structure, which consists of nt 100 – 113 of exon 12 and E12+1 – E12+53 of intron 12. The splice site lies in a 15 nt hairpin loop in the first structure and 13 × 13 nt internal loop in the second structure. U1 snRNA binding sites are part of loops in each of these structures. The V363I mutation near this splice junction, caused by a NG_007398.1:c.1087G>A substitution, is associated with primary progressive aphasia, characterized by language and speech impairment (Figure S2G).*^16^**^,^* *^18^* The result is substitution of a G-bulge with an A-bulge and an increase in ED from 5.28 to 6.39. P364S, G366R, and K369I mutations, caused by NG_007398.1:c.1090C>T, NG_007398.1:c.1096G>A, and NG_007398.1:c.1106A>T substitutions, respectively, reduce the ability of tau to promote microtubule assembly compared with wild type proteins, consistent with other intronic mutations (Figures S2H to S2J).*^19-21^* These mutations convert a GC pair to a GU pair and replace a G with an A in a hairpin loop and an A with a U in an internal loop, respectively. The mutations do not change the overall structures but increase ED was observed in each case.

A stem-loop was predicted at the intron 12-exon 13 junction with a z-score of −1.59, more than 1 SD below the average of all sequences for this junction (−0.32 ± 0.88) (Figure S1AB). The ED for this structure is 3.23.

**SUPPLEMENTARY TABLES AND FIGURES**

| Table S1: Primers used to clone fragments of tau 3′ UTR into a pmirGLO vector. | |
| --- | --- |
| Primer (nts) | Sequence (5′ to 3′) |
| 2041-2077 F | TCGAGGCTGGGGCCTCCCAAGTTTTGAAAGG-  CTTTCCTCAGCG |
| 2041-2077 R | TCGACGCTGAGGAAAGCCTTTCAAAACTTGG-  GAGGCCCCAGCC |
| 3471-3691 F | ATTACTCGAGACATTTGCTAGAGGGAGGGAG |
| 3471-3691 R | GATGTCGACGGCTTAGAGGGAAGGATGCC |
| 3695-3859 F | ATTACTCGAGTGGCACCTCTGTGCCACCTC |
| 3695-3859 R | GATGTCGACCTGCAGGTCTGTAGATGGGAC |

| Table S2: MicroRNAs predicted to bind to the tau 3′ UTR. | | |
| --- | --- | --- |
| miRNA | Recognition element in 3′ UTR (5′ to 3′) | Location of binding site in 3′ UTR |
| hsa-miR-34a | CCGTGAGAGCCCAATCACTGCCT | 1172-1194 |
| hsa-miR-132 | TGATTTACACTGACTGTT | 4104-4121 |
| hsa-miR-219 | GATCTTAAATGAGGACAATCC | 3162-3182 |
| hsa-miR-181c | GCTTTCTGTCTGTGAATGT | 4056-4074 |
| hsa-miR-485-5p | AATGTCCCGAATTCCCAGCCTCA | 1216-1238 |
| hsa-miR-642 | GAGGGTGGGGGGAGGGAC | 4019-4036 |

| **A** | **B** | **C** |
| --- | --- | --- |
| 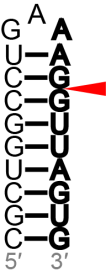 | 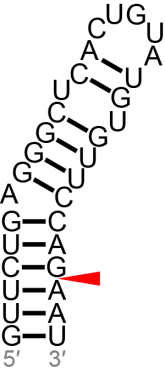 | 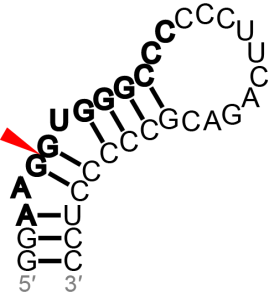 |
| Exon 1-intron 1  nt 137-150 of exon 1 and E1+1 to E1+7 of intron 1 | Intron 1-exon 2  nt E2−29 to E2−1 of I1 and 1-3 of E2 | Exon 2-intron 2  nt 82-87 of E2 and E2+1 to E2+27 of I2 |
| **D** | **E** | **F** |
| 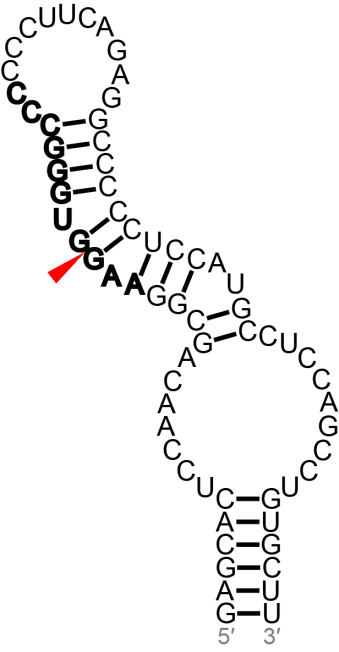 | 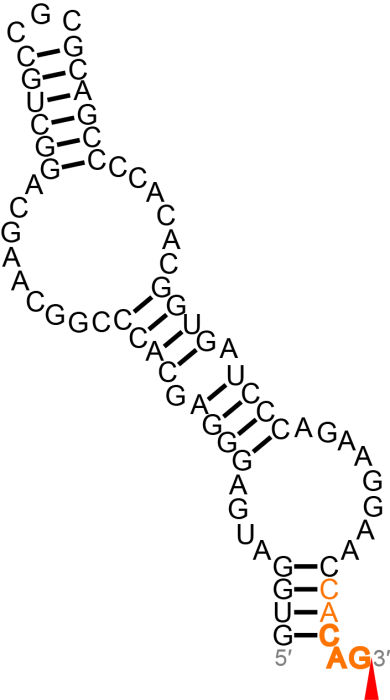 | 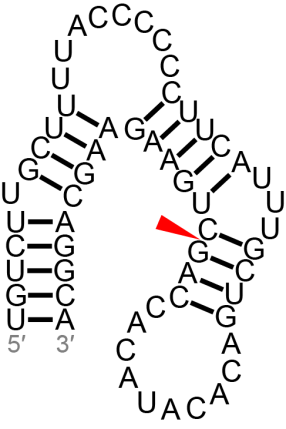 |
| Exon 2-intron 2  nt 68-87 of E2 and E2+1 to E2+46 of I2 | Exon 3-intron 3  nt 18-87 of E3 | Intron 3-exon 4  nt E4−43 to E4−1 of intron 3 and 1-15 of exon 4 |

Figure S1: Predicted structures at other exon-intron junctions. Splice sites are denoted with red arrowheads. Potential U1 snRNA binding sites are denoted by bold nucleotides.

| **G** | **H** | **I** |
| --- | --- | --- |
| 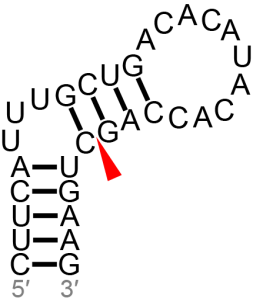 | 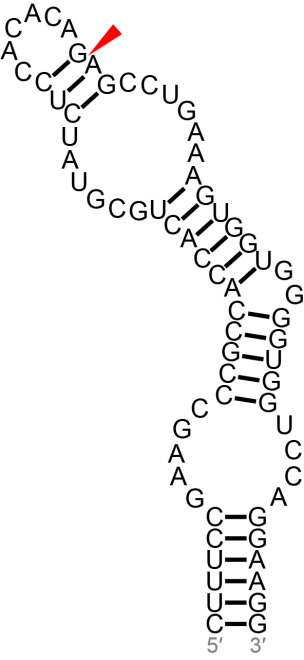 | 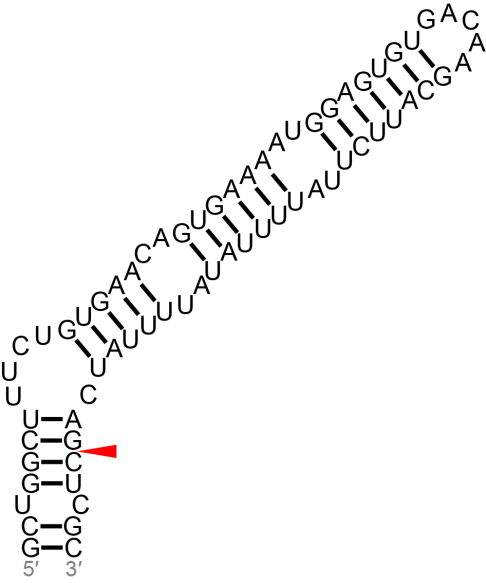 |
| Intron 3-exon 4  nt E4−25 to E4−1 of intron 3 and 1-6 of exon 4 | Intron 4-exon 4A  nt E4A−38 to E4A−1 of intron 4 and 1-31 of exon 4A | Intron 4A-exon 5  nt E5−63 to E5−1 of intron 4A and 1-5 of exon 5 |
| **J** | **K** | **L** |
| 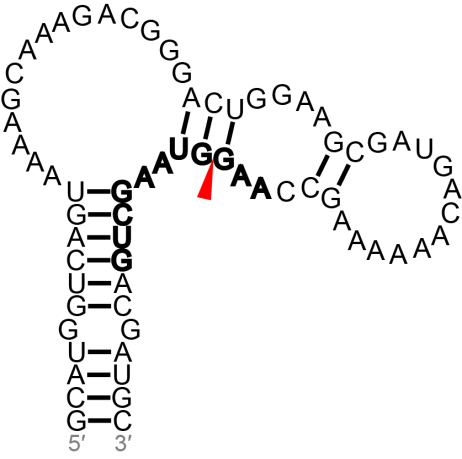 | 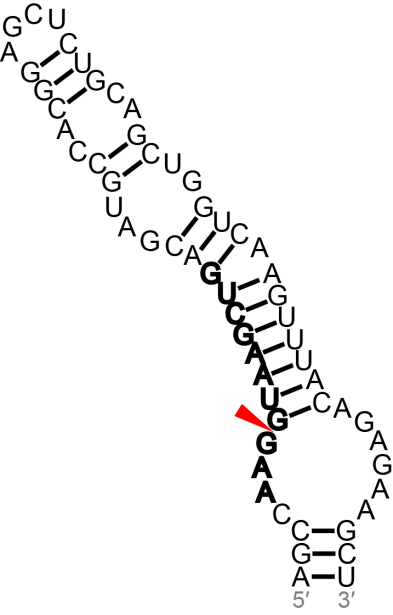 | 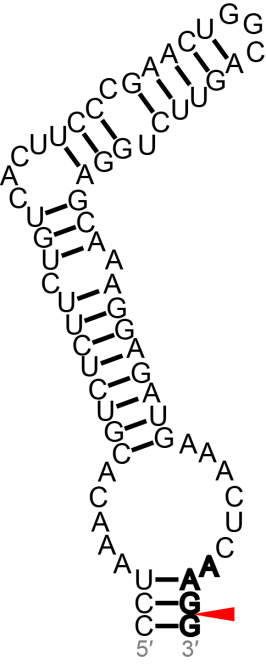 |
| Exon 5-intron 5  nt 4-56 of exon 5 and E5+1 to E5+15 of intron 5 | Exon 5-intron 5  nt 50-56 of exon 5 and E5+1 to E5+53 of intron 5 | Exon 6-intron 6  nt 133-198 of exon 6 and E6+1 of intron 6 |

Figure S1: Predicted structures at other exon-intron junctions. Splice sites are denoted with red arrowheads. Potential U1 snRNA binding sites are denoted by bold nucleotides.

| **M** | **N** | **O** |
| --- | --- | --- |
| 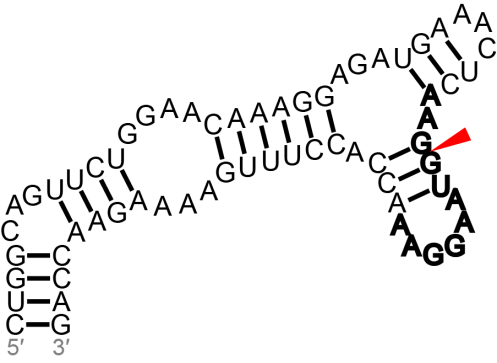 | 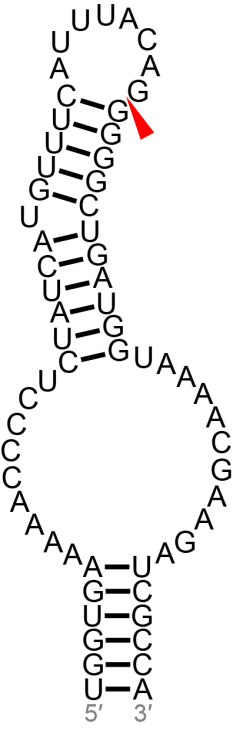 | 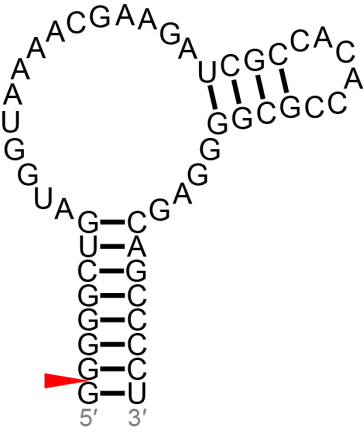 |
| Exon 6-intron 6  nts 164-198 of exon 6 and E6+1 to E6+29 of intron 6 | Intron 6-exon 7  nt E7−35 to E7−1 of intron 6 and 1-28 of exon 7 | Intron 6-exon 7  nt E7−1 of intron 6 and 1-48 of exon 7 |
| **P** | **Q** | **R** |
| 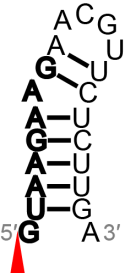 | 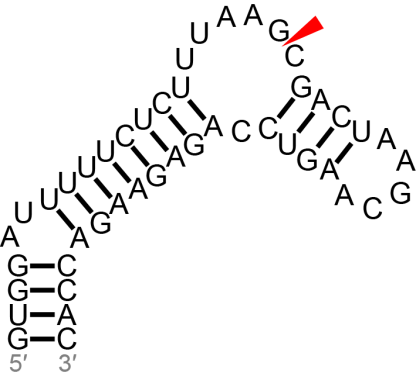 | 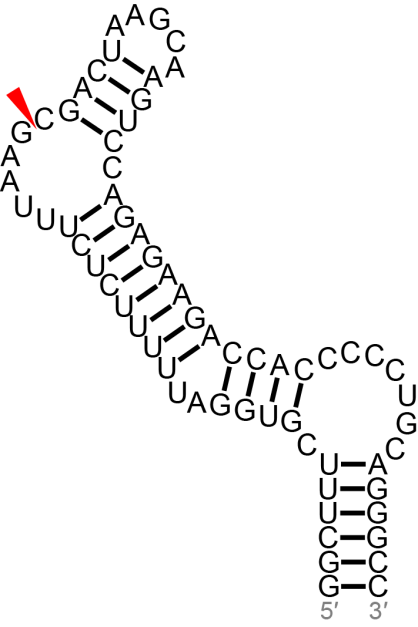 |
| Exon 7-intron 7  nt E7+1 to E7+21 of intron 7 | Intron 7-exon 8  nt E8−19 to E8−1 of intron 7 and 1-27 of exon 8 | Intron 7-exon 8  nt E8−26 to E8−1 of intron 7 and 1-40 of exon 8 |

Figure S1: Predicted structures at other exon-intron junctions. Splice sites are denoted with red arrowheads. Potential U1 snRNA binding sites are denoted by bold nucleotides.

| **S** | **T** | **U** |
| --- | --- | --- |
| 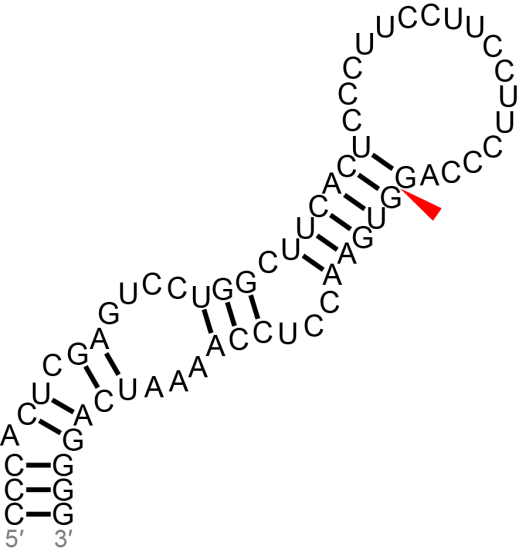 | 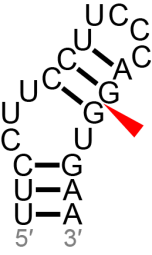 | 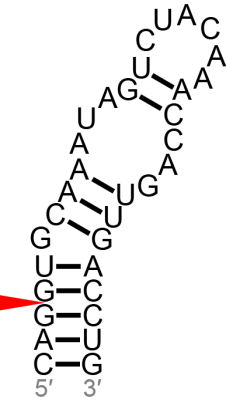 |
| Intron 8-exon 9  nt E9−41 to to E9−1 of intron 8 and 1-21 of exon 9 | Intron 8-exon 9  nt E9−15 to E9−1 of intron 8 and 1-5 of exon 9 | Intron 10-exon 11  nt E11−3 to E11−1 of intron 10 and 1-30 of exon 11 |
| **V** | **W** | **X** |
| 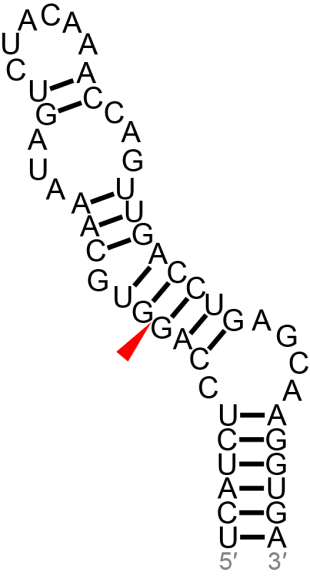 | 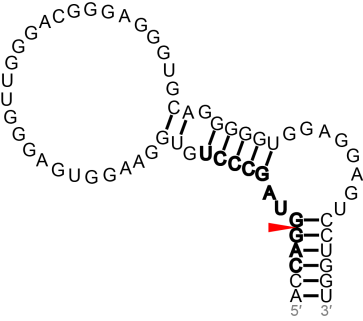 | 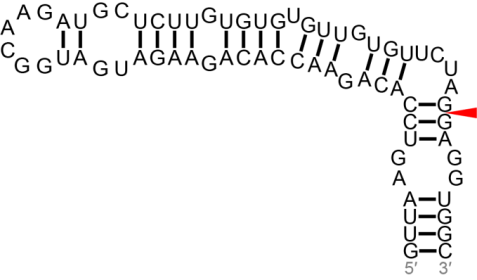 |
| Intron 10-exon 11  nt E11−10 to E11−1 of intron 10 and 1-40 of exon 11 | Exon 11-intron 11  nt 78-82 of exon 11 and E11+1 to E11+60 of intron 11 | Intron 11-exon 12  nt E12−61 to E12−1 of intron 11 and 1-8 of exon 12 |

Figure S1: Predicted structures at other exon-intron junctions. Splice sites are denoted with red arrowheads. Potential U1 snRNA binding sites are denoted by bold nucleotides.

| **Y** | **Z** |
| --- | --- |
| 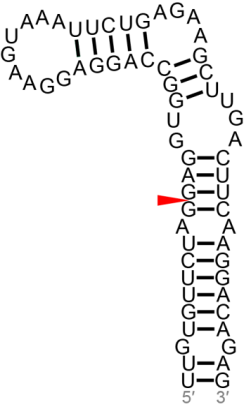 | 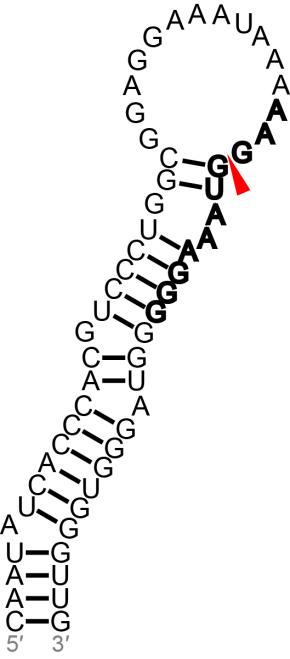 |
| Intron 11-exon 12  nt E12−11 to E12−1 of intron 11 and 1-51 of exon 12 | Exon 12-intron 12  nt 77-113 of exon 12 and E12+1 to E12+22 of intron 12 |
| **AA** | **AB** |
| 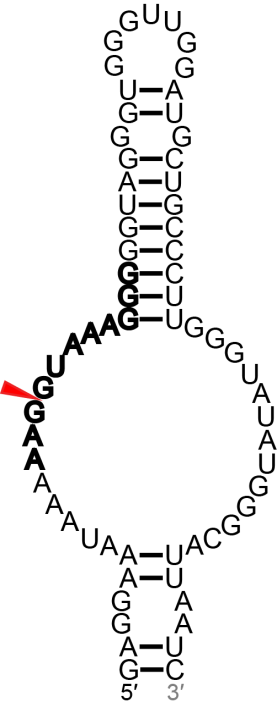 | 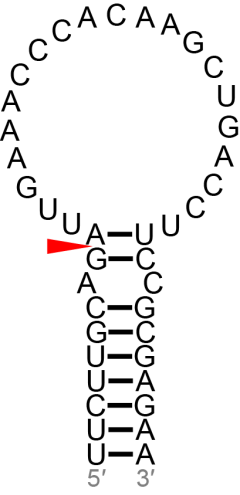 |
| Exon 12-intron 12  nt 100-113 of exon 12 and E12+1 to E12+53 of intron 12 | Intron 12-exon 13  nt E13−9 to E13−1 of intron 12 and 1-32 of exon 13 |

Figure S1: Predicted structures at other exon-intron junctions. Splice sites are denoted with red arrowheads. Potential U1 snRNA binding sites are denoted by bold nucleotides.

| **A** | **B** | **C** |
| --- | --- | --- |
| 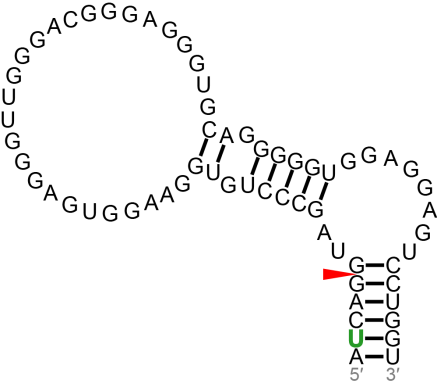 | 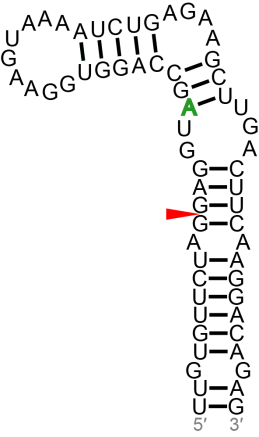 | 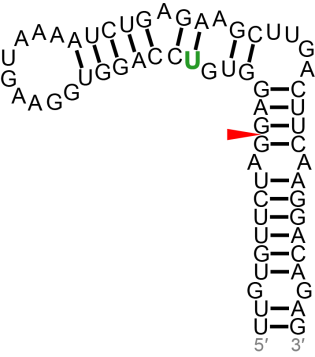 |
| Exon 11-intron 11 P332S  nt 78-82 of exon 11 and E11+1 to E11+60 of intron 11 | Intron 11-exon 12 G335S  nt E12−61 to E12−1 of intron 11 and 1-8 of exon 12 | Intron 11-exon 12 G335V  nt E12−61 to E12−1 of intron 11 and 1-8 of exon 12 |
| **D** | **E** | **F** |
| 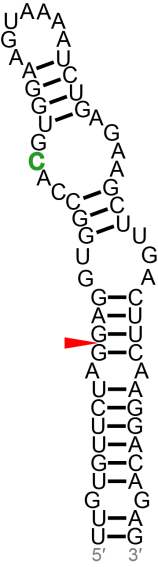 | 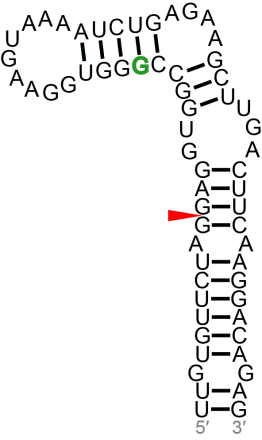 | 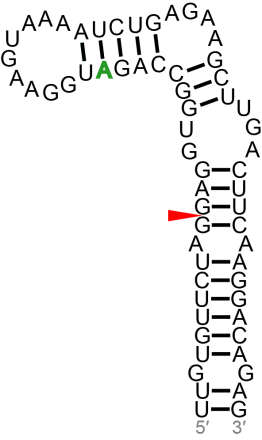 |
| Intron 11-exon 12 Q336H  nt E12−11 to E12−1 of intron 11 and 1-51 of exon 12 | Intron 11-exon 12 Q336R  nt E12−11 to E12−1 of intron 11 and 1-51 of exon 12 | Intron 11-exon 12 V337M  nt E12−11 to E12−1 of intron 11 and 1-51 of exon 12 |

Figure S2: Predicted structures at exon-intron junctions that contain mutations. Shown are structures containing mutations that only result in base pair or loop nucleotide substitutions in the wild type structure. Mutations are denoted in green. Splice sites are denoted with red arrowheads.

| **G** | **H** |
| --- | --- |
| 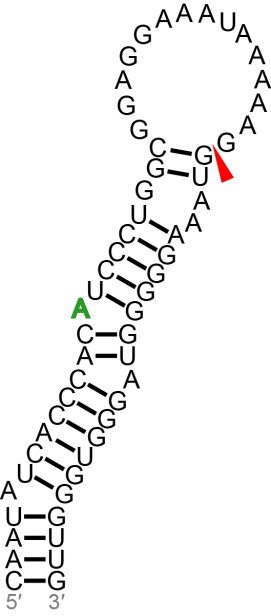 | 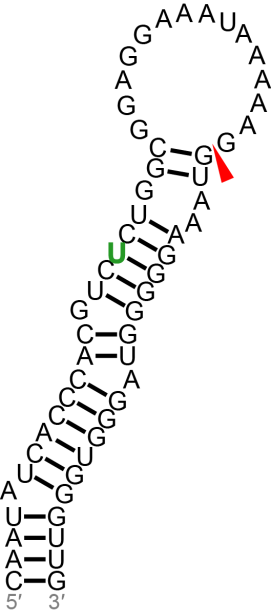 |
| Exon 12-intron 12 V363I  nt 77-113 of E12 and E12+1 to E12+22 of I12 | Exon 12-intron 12 P364S  nt 77-113 of E12 and E12+1 to E12+22 of I12 |
| **I** | **J** |
| 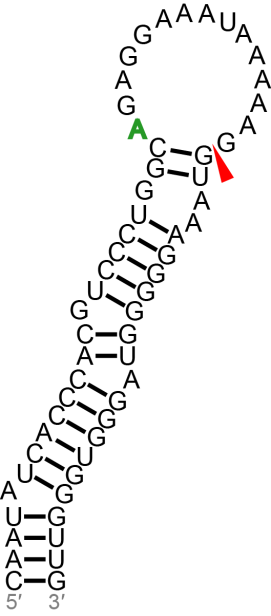 | 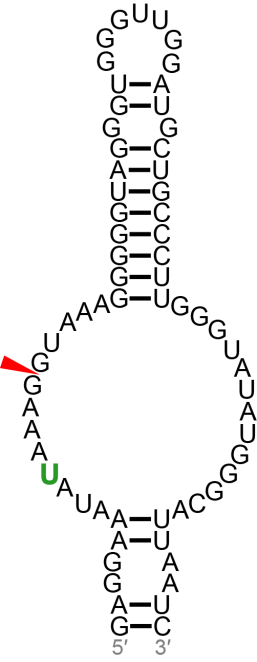 |
| Exon 12-intron 12 G366R  nt 77-113 of E12 and E12+1 to E12+22 of I12 | Exon 12-intron 12 K369I  nt 100-113 of E12 and E12+1 to E12+53 of I12 |

Figure S2: Predicted structures at exon-intron junctions that contain mutations. Shown are structures containing mutations that only result in base pair or loop nucleotide substitutions in the wild type structure. Mutations are denoted in green. Splice sites are denoted with red arrowheads.

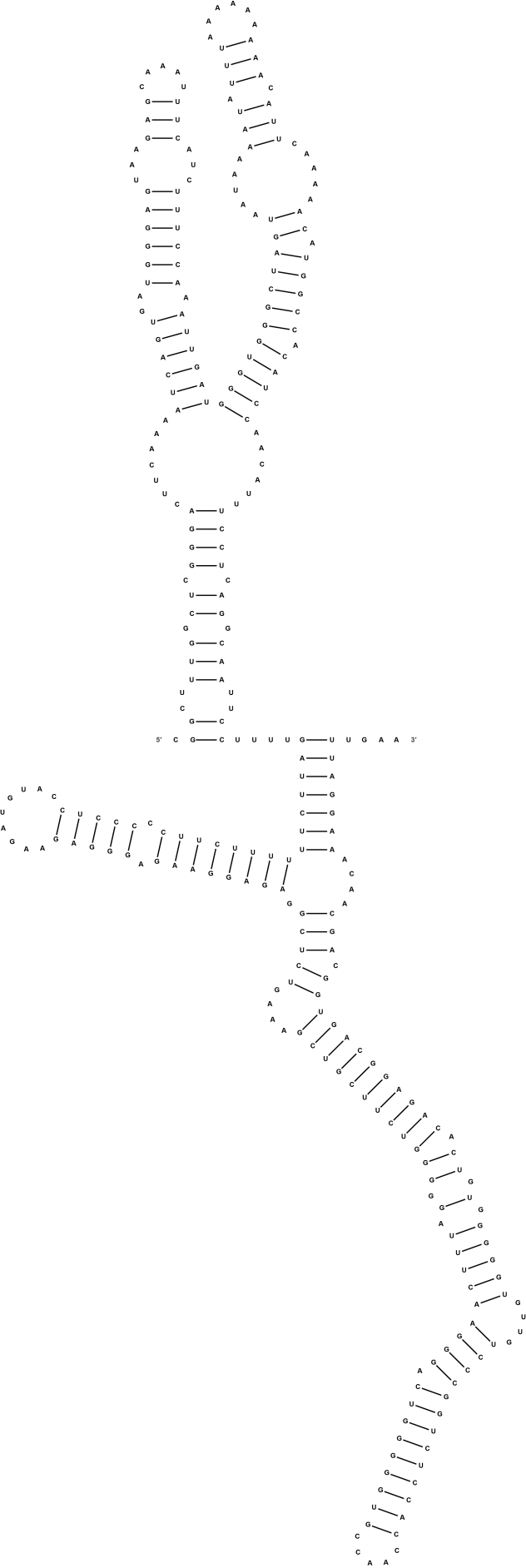

Figure S3: Positions 151-450 of the 3′ UTR.

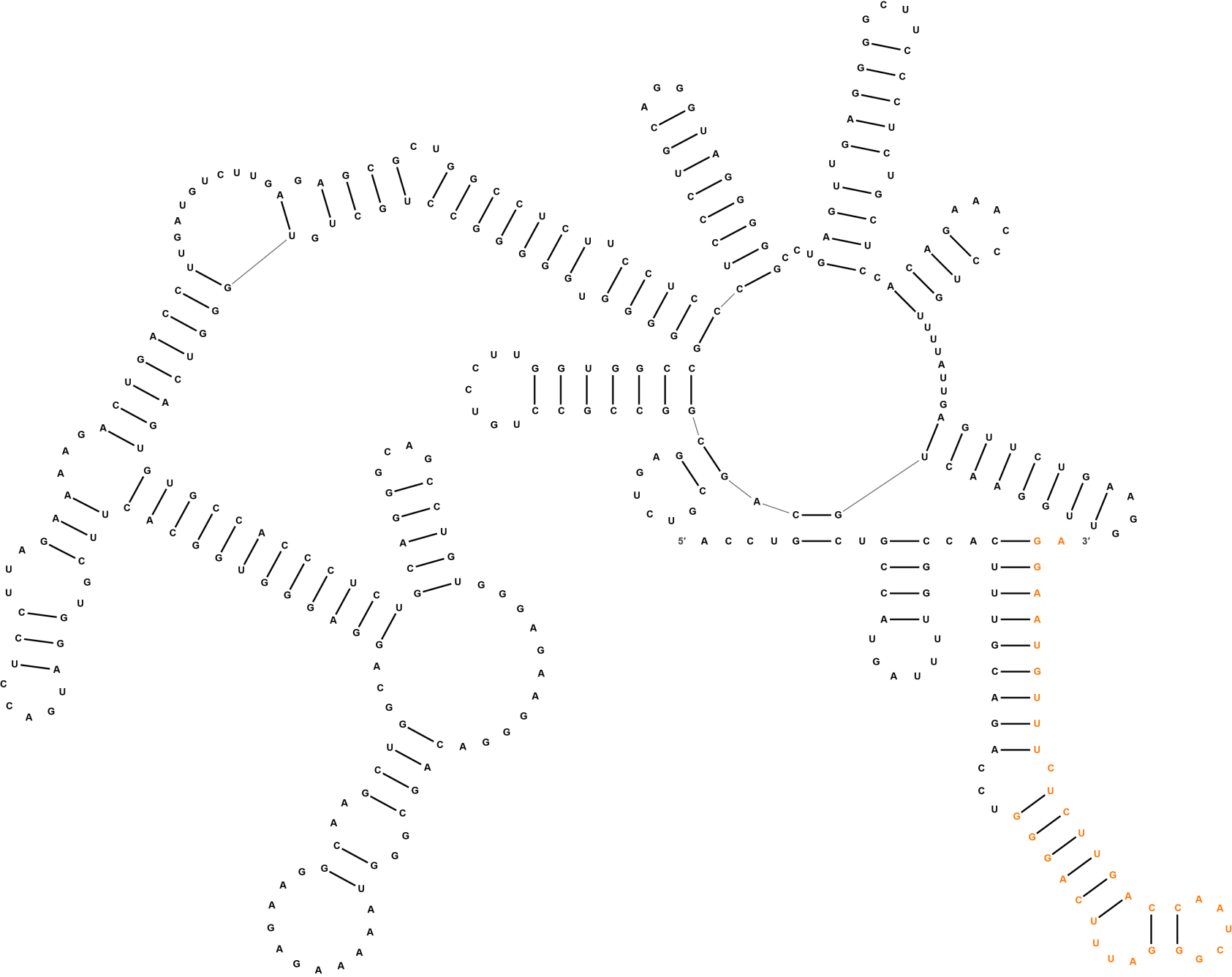

Figure S4: Positions 661-1020 of the 3′ UTR. Deletion of residues labeled orange from a luciferase reporter construct increased luciferase activity in transfected cells compared to those expressing a wild type 3′ UTR construct.

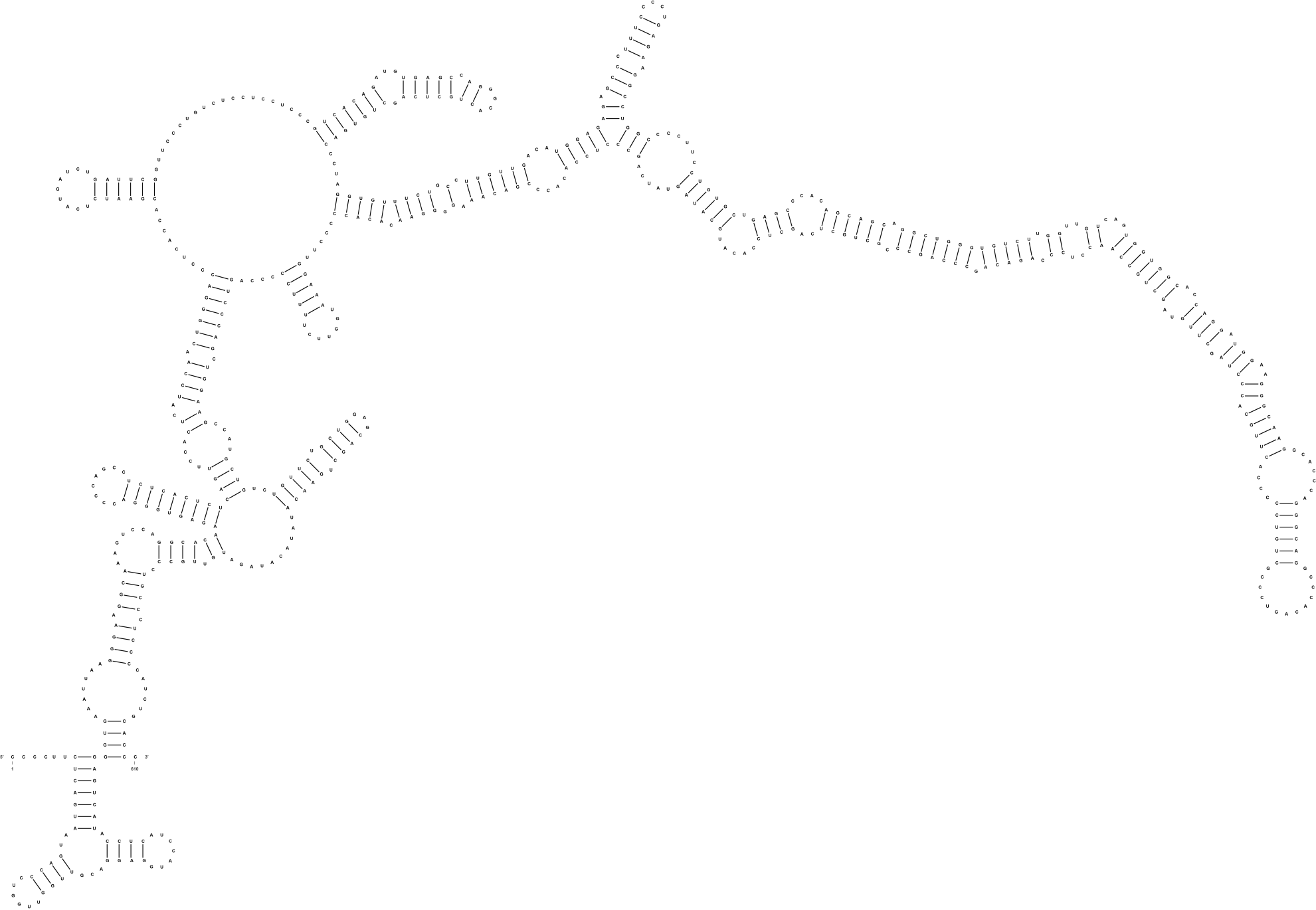

Figure S5: Positions 1241-1850 of the 3′ UTR.

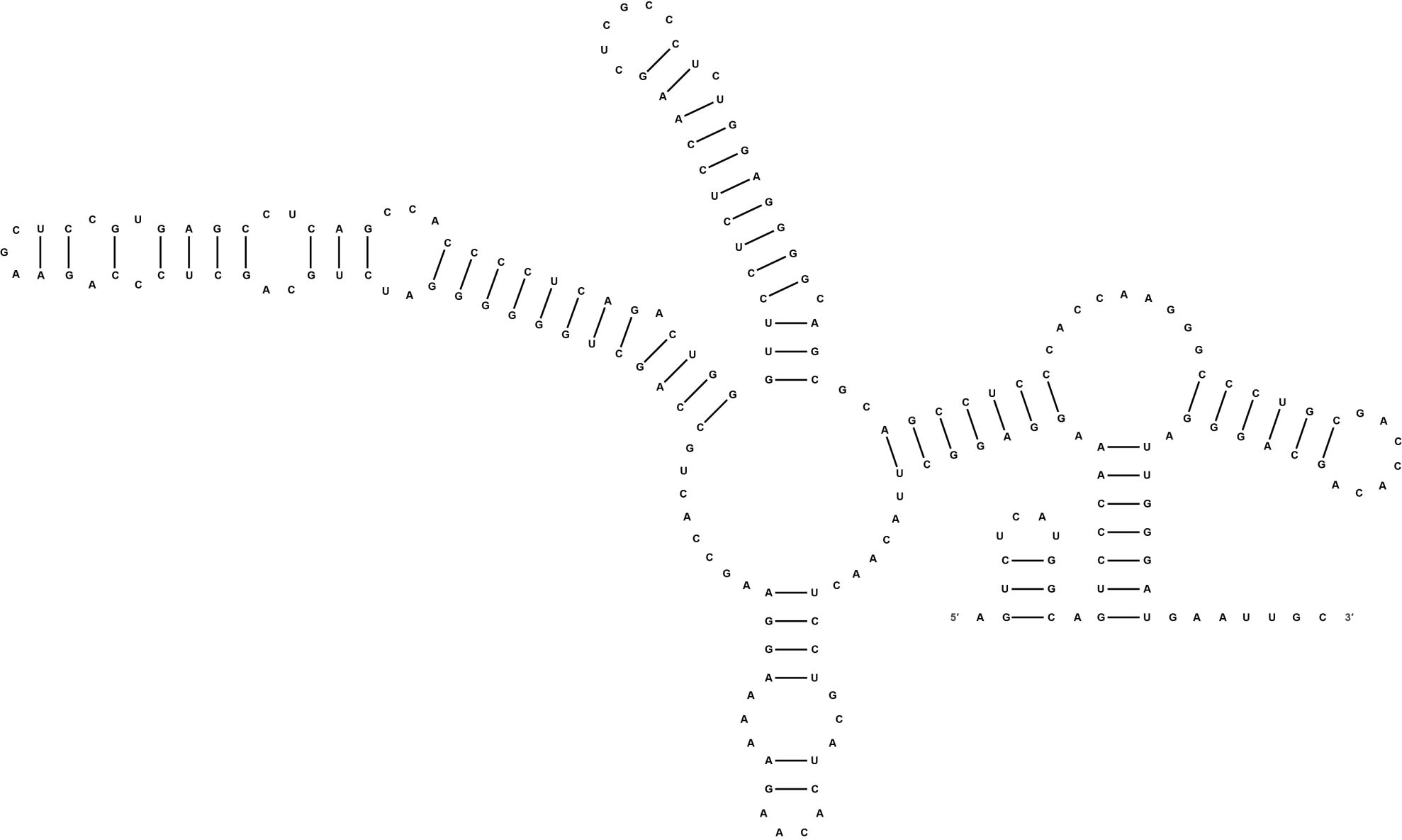

Figure S6: Positions 2281-2490 of the 3′ UTR.

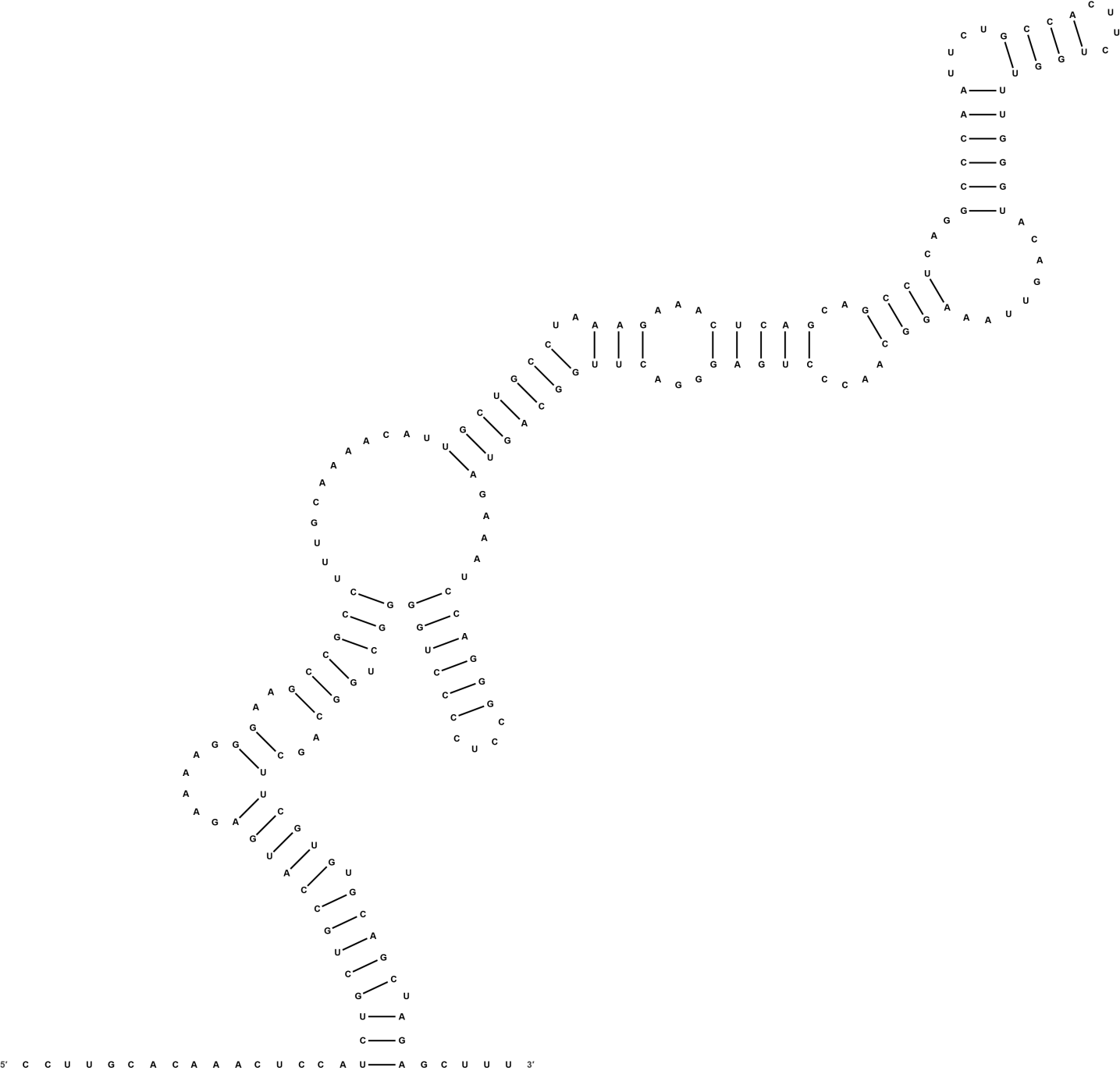

Figure S7: Positions 2601-2800 of the 3′ UTR.

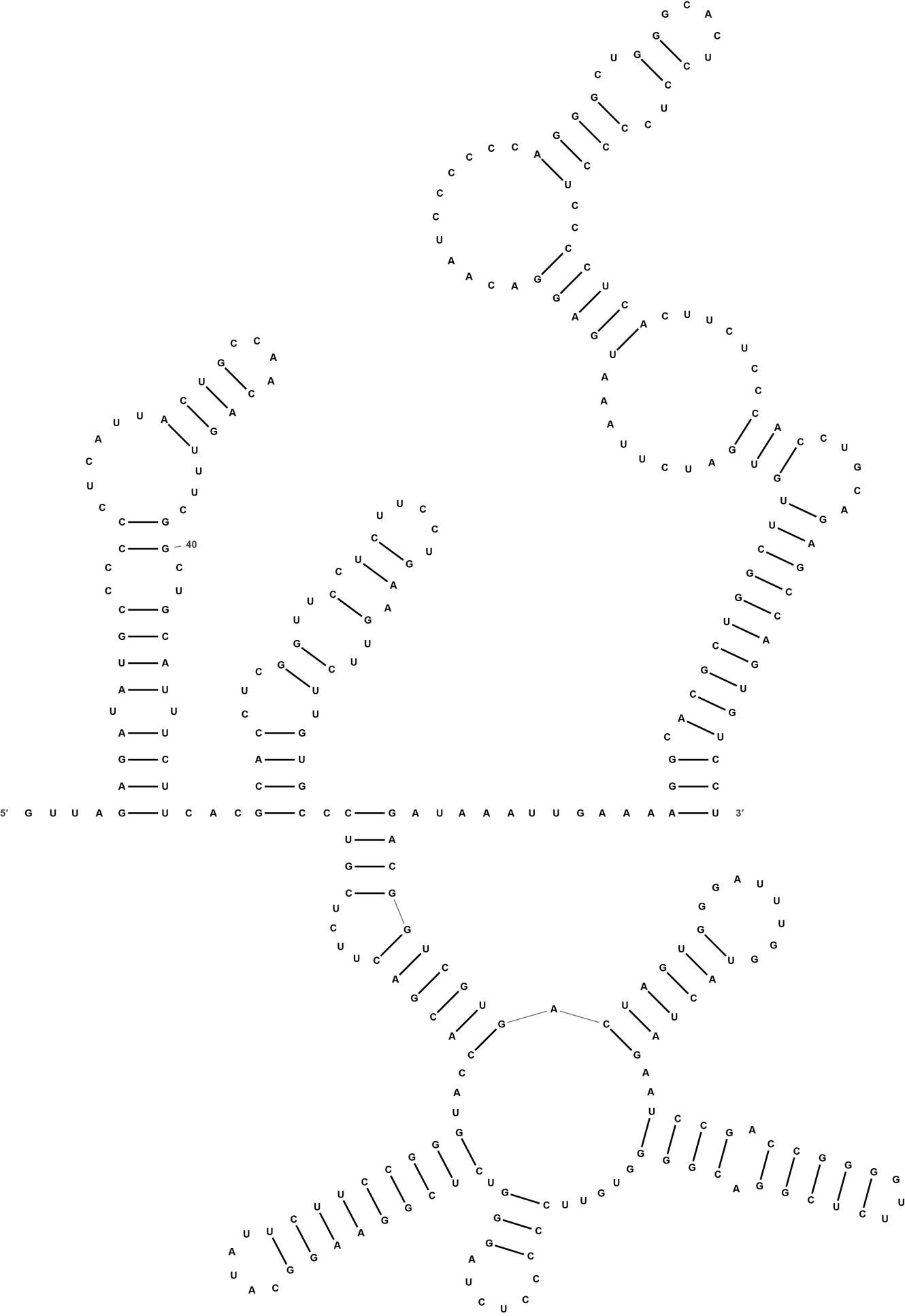

Figure S8: Positions 2931-3240 of the 3′ UTR.
